# Centromere chromatin may regulate the Flip-flop type of the Mating Type Switching in methylotrophic yeast *Ogataea polymorpha*

**DOI:** 10.1101/2025.09.09.675088

**Authors:** Nodoka Fukuyama, Kaoru Takegawa, Hiromi Maekawa

**Affiliations:** Laboratory of Applied Microbiology, Department of Bioscience and Biotechnology, Faculty of Agriculture, Graduate School of Bioresource and Bioenvironmental Sciences, Kyushu University, 744 Motooka, Nishi-ku, Fukuoka 819-0395, Japan; Laboratory of Fungal Cell Biology, Faculty of Agriculture, Graduate School of Bioresource and Bioenvironmental Sciences, Kyushu University, 744 Motooka, Nishi-ku, Fukuoka 819-0395, Japan

## Abstract

Centromere chromatin has been implicated in silencing the mating type (*MAT*) locus, which resides within a chromosomal region occupied by OpCse4, the CENP-A homolog, in the Flip-flop type of Mating Type Switching (MTS) in the methylotrophic yeast *Ogataea polymorpha*. In this study, we investigated the role of centromere chromatin in the recombination event between *MAT*-adjacent inverted repeats (IRs) that drives MTS. Our results demonstrate that the position of the centromere-proximal IR is critical for efficient recombination between the IRs. Mutants lacking functional OpCse4 or its chaperone OpScm3 exhibit MTS-deficient phenotypes, supporting an active role for centromere chromatin in facilitating MTS. Additionally, we identified OpRad6 and OpBre1 as essential for MTS, and found that deletion of the C-terminal acidic tail of OpRad6 alone is sufficient to disrupt switching. This suggests that monoubiquitylated histone H2B may contribute to MTS either directly or indirectly. Collectively, our findings highlight a direct and functional involvement of centromere chromatin in promoting Flip-flop type MTS in *O. polymorpha*.

## Introduction

Homothallic mating capability is observed in many Ascomycetous yeasts. Evolutionary transitions from heterothallism to homothallism have been reported in most of the major clades of budding yeasts(Krassowski et al. 2019). Homothallism has two major forms: In primary homothallism, which consists of two mating type (MAT) loci, cells are presumed to mate with any other cell, although the details have not been investigated. In secondary homothallism, each cell expresses a mating type identity at a time and can switch mating type. Two types of Mating Type Switching (MTS) mechanisms are known in budding yeasts: the 3MAT type and the Flip-flop type. The 3MAT type MTS is identified only in Saccharomycetaceae in budding yeasts. The mechanism has been well studied in *Saccharomyces cerevisiae* (Lee and Haber 2015; Wolfe and Butler 2022). Two of three copies of mating type genes at *HML* and *HMR* are maintained silent through heterochromatin formed by the Sir complex, and introduction of a site-specific double-strand break DNA at the active *MAT* locus initiates the MTS, and repair of the DSB by the Synthesis-Dependent Strand Annealing (SDSA) pathway using a silent *HML* or *HMR* as a donor, results in a gene conversion at the *MAT* locus. The Flip-flop type consists of two MAT loci on the same chromosome and their adjacent repeat sequences, first reported in methylotrophic yeasts *Ogataea polymorpha* and *Komagataella phaffii* (Maekawa and Kaneko 2014; Hanson, Byrne, and Wolfe 2014). Distinct from the 3MAT system, the molecular mechanism of the Flip-flop type is presumed to be homologous recombination with a crossover (CO) outcome at the inverted repeats (IRs) adjacent to the *MAT*s, resulting in the swapping of the chromosomal locations of two MATs (Maekawa and Kaneko 2014; Hanson, Byrne, and Wolfe 2017, 2014).

*O. polymorpha* genome carries two *MAT*s, a *MAT****a*** and a *MATα*, close to the centromere on chromosome 4, that are sandwiched between inverted repeats (IRs). Only the centromere-proximal *MAT* locus is occupied by the CENP-A homologue and kept silent, which is essential for a cell to behave as haploid and express the mating type identity of the centromere-distal *MAT* locus (Hanson, Byrne, and Wolfe 2014; Yamamoto et al. 2017). The MTS is suppressed in cycling cells and occurs only after several hours of incubation under starvation conditions, around the time when cells cease mitotic divisions (Yamamoto et al. 2017; Feng et al. 2020).

Centromeres are characterised as regions that consist of nucleosomes containing CENP-A, a centromere-specific histone H3 variant, and heterochromatin in most eukaryotes. *S. cerevisiae* is an exception and has uniquely evolved point centromeres that are built on specific sequences and have 125 bp in length, and the surrounding genes are not silenced. Centromeres in most other budding yeasts are epigenetically regulated, not by DNA sequences, and are longer (from a few kb to ∼20 kb) with diverse sequences and structures (Guin, Sreekumar, and Sanyal 2020). Most budding yeasts do not have histone H3K9me or H3K27me modification which marks for heterochromatin in eukaryotes(Obuse and Nakayama 2025). Despite the lack of normal heterochromatinizing machinery, centromere/pericentromere in *Candida albicans* has heterochromatin-like characteristics and exhibit the ability to silence genes (Sreekumar et al. 2019; Freire-Benéitez, Price, and Buscaino 2016; Ketel et al. 2009), and in *O. polymorpha*, centromeric regions are transcriptionally silent, including the mating type genes at the centromere-proximal *MAT* locus (Hanson, Byrne, and Wolfe 2014). It is assumed that Cse4-containing chromatin has heterochromatin-like properties and can silence genes. Whether chromatin statuses at or sorrounding the two IRs contribute to promoting the MTS in *O. polymorpha* has not been investigated.

In this study, we analysed mutants in which the position of an IR is altered and found that the position of the centromere-proximal IR and Cse4 chromatin have active roles in the MTS. Further, we provide evidence to suggest that histone H2 monoubiquitination may contribute to the MTS mechanism. Our study shed light on the role of chromatin regulation as a fundamental component of the Flip-flop type MTS mechanism.

## Materials and Methods

### Yeast strains and plasmids

Strains and plasmids used in this study are listed in Supplementary Table S1. All yeast strains were derived from CBS4329 or NCYC495. and were generated by PCR-based methods(Janke et al. 2004). Sequences of primers used in this study are listed in Supplementary Table S2. O. polymorpha cells were transformed by electroporation. (Kiel et al. 2007; Maekawa and Kaneko 2014). To construct *iAID-CSE4* strain, *OsTIR*-*CSE4*(aa 1-53)-mAID-*CSE4* (aa 54-end)-*OpURA3* fragment was first amplified with S2-TIR9myc_Rv and HpURA3_7 using pHM1217 as template.

Then, the amplified fragment was used as a template for PCR with HpCNP1-S1 and HpCNP1-S2 to obtain the cassette fragment for yeast transformation. For *iAID-SCM3* strain, the DNA fragment for yeast transformation was amplified by PCR with OpSCM3_S1_AgTEF1 and OpSCM3_9 using pHM1219 as template. Plasmid maps and sequences for pHM1092, pHM1217, and pHM1219 are shown in Supplementary Figure S1.

### Yeast growth conditions and general methods

Yeast strains were grown either in Yeast Extract–Peptone–Dextrose (YPD) medium (1% Yeast extract, 2% Peptone, 2% Dextrose) containing 200 mg/L adenine, leucine, and uracil (YPDS) (Sherman 1991). All experiments were performed at 30 °C unless otherwise indicated. Growing cells in YPDS at 30 °C was transferred to NaKG medium, and incubated at 30 °C for the indicated hours in the Figure legends (-N samples) (Hanson, Byrne, and Wolfe 2014).

For iAID system, IAA (Merck KGaA, Darmstadt, Germany) was dissolved in ethanol to make a 500 mM stock solution, and Doxycycline (Takara Bio Inc., Shiga, Japan) was dissolved in H2O at 10 mg/mL, and stock solutions were stored at -20°C. To induce degradation of the protein fused with mAID, IAA and Doxycycline were added directly to the medium at 500 µM and 10 µg/mL, respectively, 2.5 hrs after shifting to NaKG medium.

### Determining the orientation of the region between two IRs

Orientation of the MAT-containing region was determined by PCR as previously described with the primers listed in Supplementary Table S2 (Maekawa and Kaneko 2014).

## Results and Discussions

### The IR sequences cannot promote an inversion in chromosomal arm region

We constructed a yeast strain carrying two additional IR sequences inserted into *OpADE12* locus in the same orientation as in the *MAT*-containing region (Fig.1). The orientation of the IRs was stable in YPDS medium, and an inversion did not occur after incubating in NaKG medium (Fig.1A, B), while inversion between IRs adjacent MATs occurred (Fig.1C, D). Even though the environment surrounding the IRs unit at *OpADE12* locus is different from the IRs adjacent MATs, such as the distance between two IR, locating within a coding gene, in addition to being in the non-centromeric euchromatin, the result did not disagree with the hypothesis of contribution of centromeric chromatin in the MTS.

**Figure 1.**
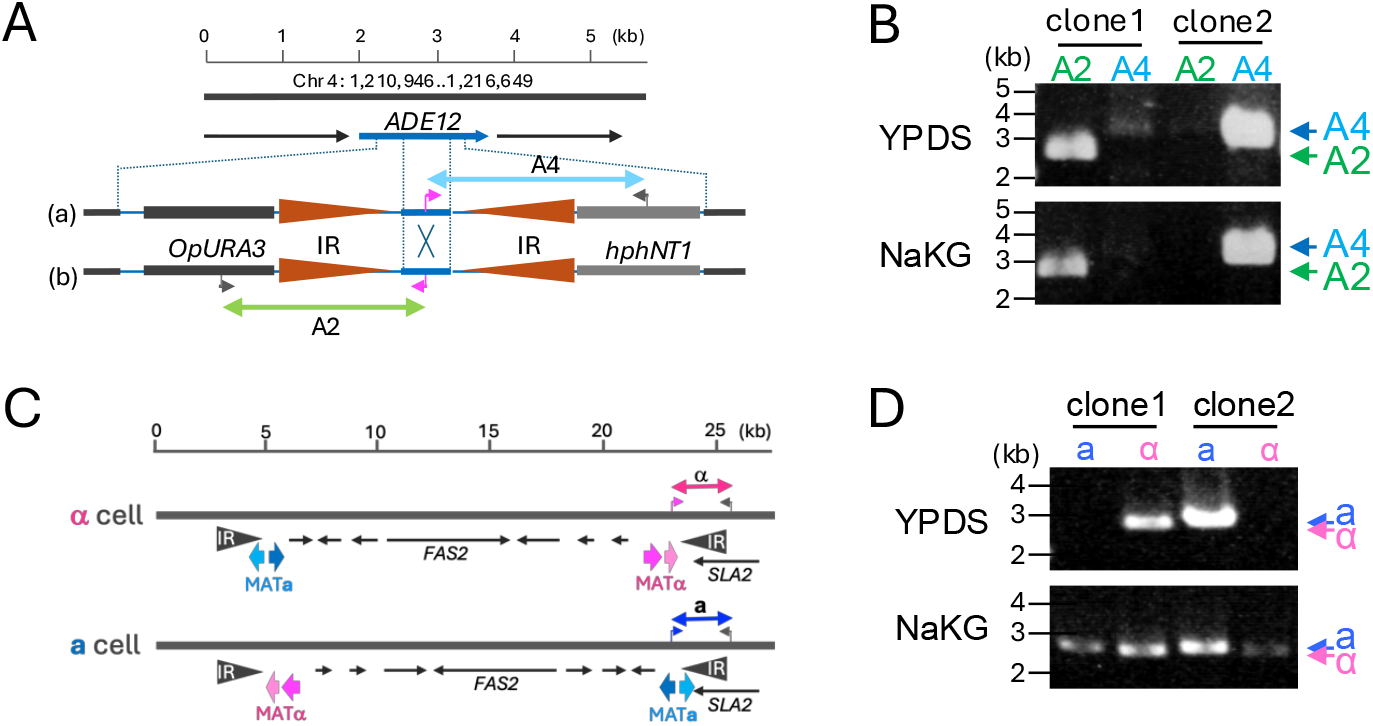
Recombination does not occur between IRs inserted into *OpADE12* locus. (A) Schematics of IR insertion into *OpADE12* locus. Double arrows indicate the PCR fragments with A2 and A4 primer sets. (B) Two independent clones of the strain carrying *Opade12::IRs* grown in YPDS were transferred into NaKG. PCR was performed to determine orientation of the IR-containing region using primer sets A2 and A4. Inversion was not induced in NaKG. (C) Schematics of MAT-containing region in **a** cell and α cell. Double arrows indicate the PCR fragments with a and α primer sets. (D) Orientation of the MAT-containing region was examined in (B).

### Position of the centromere proximal IR is crucial for promoting the MTS

The centromere-distal IR (IR1) is in the euchromatic region, while the centromere-proximal IR (IR2) is in the Cse4 (CENP-A homologue)-occupied centromeric chromatin, whose function is presumed to contribute for silencing the Cse4-occupied MAT locus (Hanson, Byrne, and Wolfe 2014). We considered the possibility that having an IR in one of these or both chromatin statuses is required for promoting recombination between the IRs. We constructed IR-position mutants in which one of the IRs is removed from the original location and placed in the middle of the IRs (Fig.2A, IR1-position mutant and IR2-position1 mutant). To prevent the expression of both *MAT*s in the centromere-proximal and distal loci, which is expected to mimic a diploid state and inhibit the inversion between the IRs, *MAT****a****1* gene was deleted in the IR2-position mutant as well as the wild-type control cells (Fig.2B, C). Inversion between IRs occurred under the starvation condition in the IR1-position mutant, but not in the IR2-position1 mutant, suggesting the location of the IR2 is important. It is possible that the IR2 placed in euchromatin prevented the inversion. Alternatively, the alteration of DNA sequence surrounding the IR2 did not allow the inversion. In order to distinguish these possibilities, 7.8 kb marker fragment of 3mCherry-*ScLEU2*-*natNT2* was inserted at the centromere-proximal end of the IR2, which would move the IR towards the arm region without altering the DNA sequence of the centromere-distal side of IR2 (IR2-position2 mutant) (Fig.2A, D).

**Figure 2.**
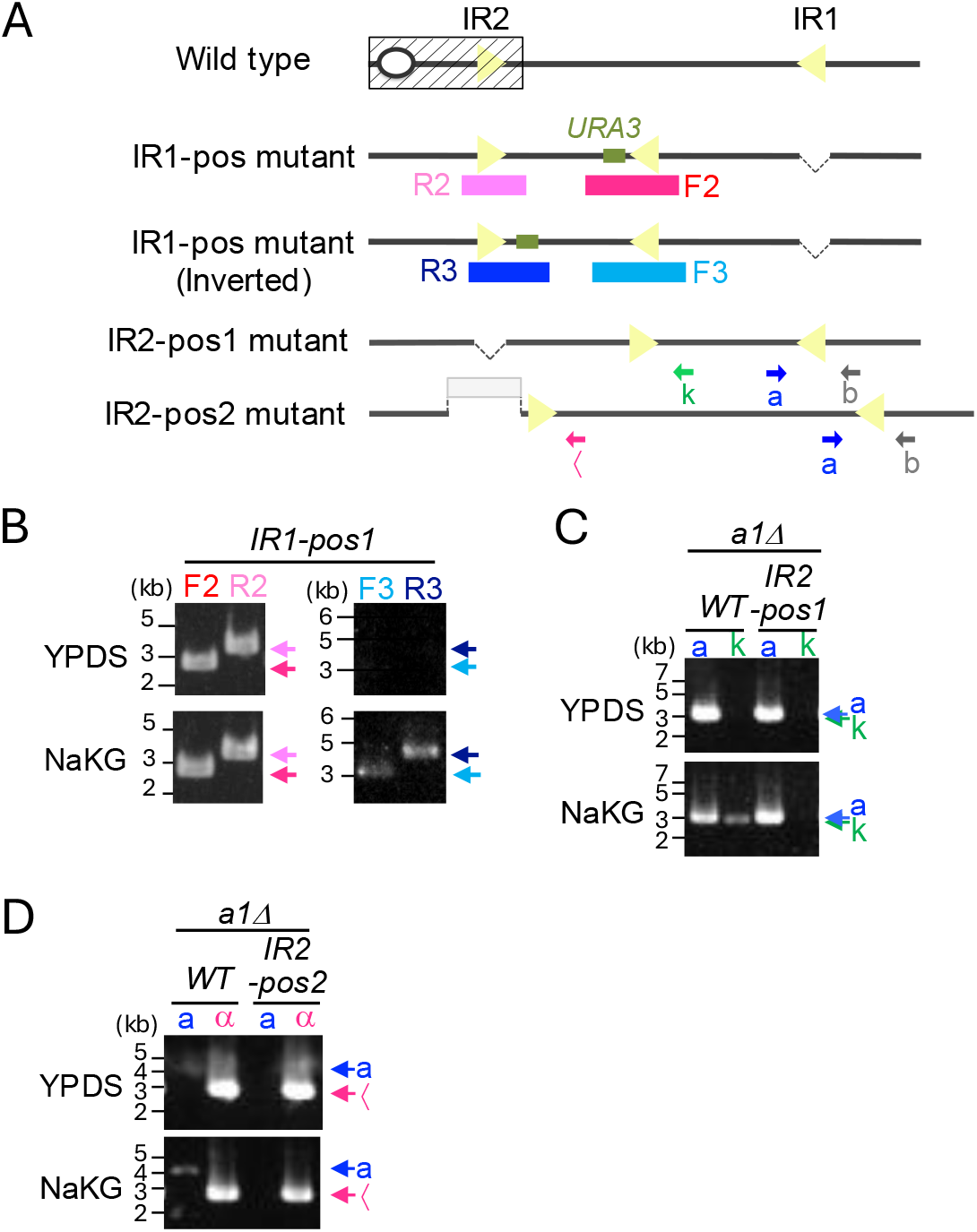
Location of the centromere-proximal IR is important for promoting the MTA. (A) Schematics of IR position mutants. Coloured boxes indicate PCR fragments with F2, F3, R2, and R3 primer sets. Blue, magenta, green, and grey arrows indicate the annealing sites of primers a, α, and k. PCRs were performed with primer b and primer a, α, or k. (B) IR pos1 mutant grown in YPDS were transferred into NaKG. Inverted allele was detected in F3 and R3 PCRs in NaKG condition. (C) (D) Wild type and IR2 position mutants with *a1Δ* background were grown in YPDS and transferred into NaKG. PCR was performed to determine orientation of the MAT-containing region using primer a, α, or k together with primer b. Inverted allele was not detected in the IR2 position mutants.

The IR-inversion was observed in ***a****1Δ* cells, albeit somewhat weakly. In contrast, the IR2-position2 mutant cells did not show a sign of inversion after incubating in NaKG medium. These results indicated that an IR needs to reside in the centromeric chromatin for promoting the MTS.

### Centromere chromatin is required for the MTS

If Cse4-containing chromatin is essential for the MTS, mutants for *OpCSE4* or *OpSCM3*, a Cse4p specific chaperone, would exhibit defects or reduction of the MTS. As deletion mutants for *OpCSE4* and *OpSCM3* would be inviable, we constructed conditional mutants using the improved auxin-inducible degron system (iAID) (Tanaka et al. 2015; Maekawa et al. 2022) (Fig. 3A). Logarithmically growing cells in YPD medium were transferred to the NaKG starvation medium, and incubated for 2 hours to allow the initial starvation response to occur, and then 10 µg/mL doxycycline and 500 µM IAA were added to repress the transcription of the AID-fused gene and to activate the degradation of the AID-tagged proteins. The inverted orientation was not observed in *iAID-CSE4* and *iAID-SCM3* cells (Fig. 3B, C). To eliminate a possibility that reduction of OpCse4 or OpScm3 compromise the silencing the centromere-proximal *MAT* and both **a** and α mating type genes are expressed, mimicking the diploid state, the experiment was performed in cells carrying ***a****1Δ* (Fig. 3D). Although the MTS in ***a****1Δ* cells was weaker than wild type cells, it was further reduced to undetectable level in *iAID-CSE4* and *iAID-SCM3* cells. These results indicated the active involvement of centromeric chromatin in the IR-inversion.

**Figure 3.**
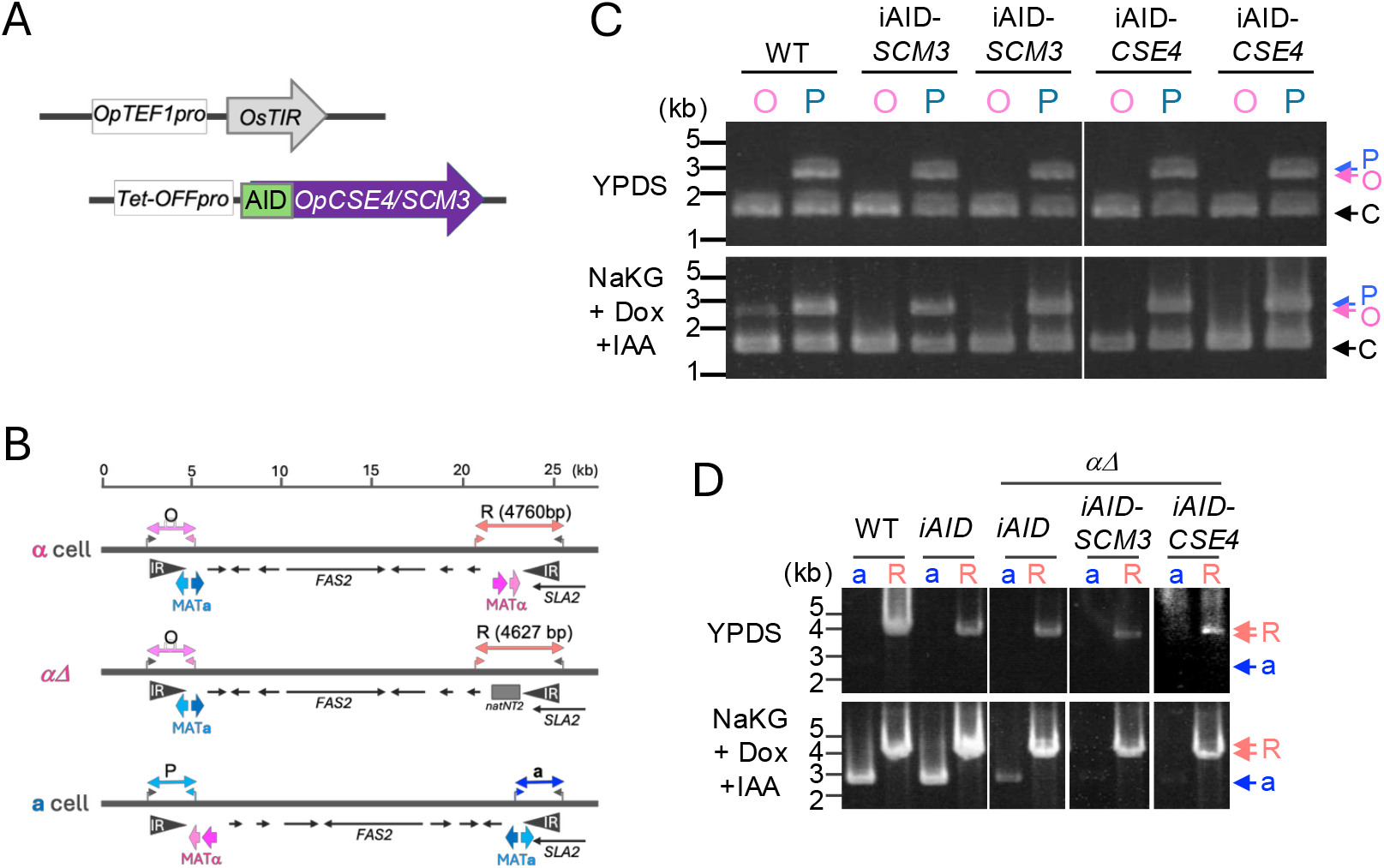
The MTS does not occur in centromere nucleosome mutants. (A) Schematics of iAID-CNP1 and iAID-SCM3. (B) Primer sets a, O, P, and R are indicated as double arrows. (C) Wild type, iAID-SCM3, iAID-CNP1 cells were grown in YPDS, then transferred into NaKG. Doxycycline and IAA were added after 2.5 hours incubation in NaKG. PCR was performed to determine orientation of the MAT-containing region using primer sets O and P. (D) Cells with indicated genotype were grown in YPDS and transferred into NaKG. PCR was performed to determine orientation of the MAT-containing region using primer sets a and R.

### OpRad6 together with OpBre1 plays a pivotal role in the MTS

The above results indicated the chromatin state of IRs influence the MTS. In our search for genes involved in recombination or repair, we noticed that the MTS was defective in a deletion mutant for *OpRAD6*, homologue of *S. cerevisiae RAD6* (Fig.4A). Rad6 is a highly conserved E2 ubiquitin conjugation enzyme among eukaryotes, and is known to be multifunctional, participating in several pathways including ubiquitin-mediated protein degradation, DNA repair, and post-replication repair pathways. In *S. cerevisiae*, Rad6 forms protein complexes with distinct E3 ubiquitin ligases and is involved in several cellular pathways. Rad6, in a complex with Rad18, monoubiquitinate lysine 167 (K167) of PCNA and is involved in postreplication DNA repair (Hoege et al. 2002; Jentsch, McGrath, and Varshavsky 1987). It also forms a complex with Bre1 and monoubiquinates lysine 123 (K123) on histone H2B for epigenetic transcriptional activation of a specific set of genes, as well as chromatin regulation through influencing other histone modifications, such as methylation of histone H3 at lysine 4 and 79 (Sun and Allis 2002; Kao et al. 2004; Xiao et al. 2005; Deng et al. 2020). It is also involved in N-end rule-dependent protein degradation in association with Ubr1 (Dohmen et al. 1991; Sung et al. 1991; Watkins et al. 1993). To clarify which function is required for the MTS, we analysed deletion mutants for *OpRAD18* and *OpBRE1*. The MTS was deficient in *Opbre1Δ*, but not in *Oprad18Δ* cells, suggesting that OpRad6-OpBre1 functions in the MTS, while OpRad6-OpRad18 is dispensable (Fig.4B). To further confirm, we constructed a *Oprad6* mutant in the OpBre1-interacting domain. In *S. cerevisiae*, ScRad6 interacts with ScBre1 through the acidic tail at the C-terminus(Yadav et al. 2024; Deng et al. 2020). Amino acid alignments of Rad6 from budding yeast species revealed that the acidic tail is conserved in *O. polymorpha*, but not in early-diverged budding yeast species such as *Y. lipolytica*, the fission yeast *Schizosaccharomyces pombe*, or other eukaryotes (Fig.5A). *OpRAD6* gene fused with GFP at the C-terminus was fully functional in the MTS, but the MTS was almost abolished in cells with deletion of the C-terminus acidic tail (aa150-183) (*rad6ΔC-GFP*) (Fig. 5B). Interestingly, fusing with a highly acidic 5xflag tag, instead of GFP, restored the MTS function in *rad6ΔC* cells (Fig. 5C), highlighting the role of an acidic domain at C-terminus. These results indicated that OpRad6-OpBre1 complex plays a pivotal role in promoting the MTS, likely through ubiquitinating histone H2B. Whether the OpRad6-OpBre1 complex functions through transcriptional activation of genes required for the MTS, or it functions by modulating chromatin at or near the IRs and *MAT*s is currently unknown. Further investigation to identify their ubiquitination targets would be necessary to clarify this point.

**Figure 4.**
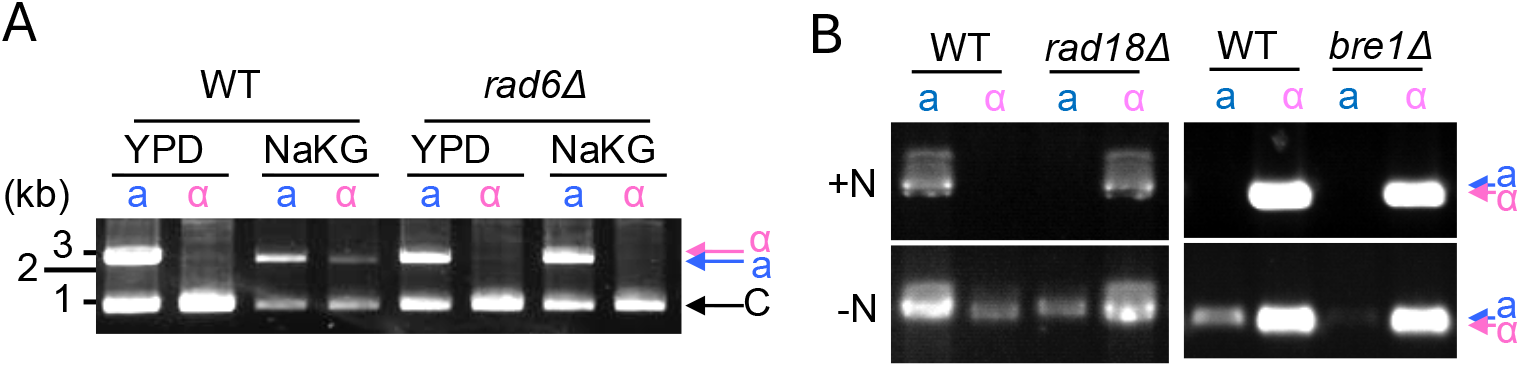
Rad6 and Bre1 contribute to the MTS. (A) (B) Cells with indicated genotype were grown in YPDS and transferred into NaKG. PCR was performed to determine orientation of the MAT-containing region using primer sets a and α indicated in Fig 1C.

**Figure 5.**
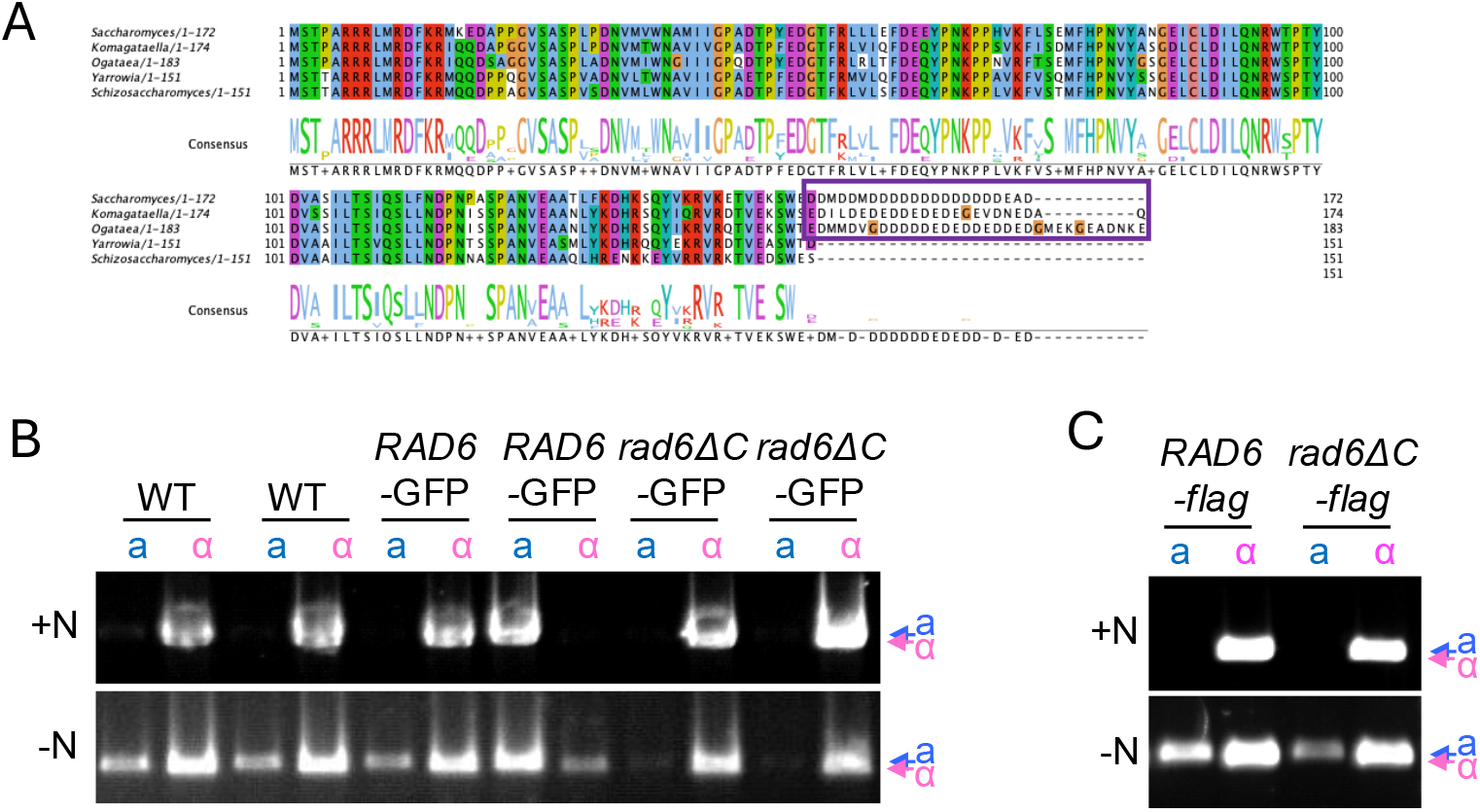
Deletion of the C-terminal acidic tail of Rad6 causes a defect in the MTS. (A) Alignment of Rad6 homologues in *S. cerevisiae, Komagataella phaffii, O. polymorpha, Yarrowia lipolytica*, and the fission yast *Schizosaccharomyces pombe*. Purple box indicates C-terminal acidic tail. (B) (C) Cells with indicated genotype were grown in YPDS and transferred into NaKG. PCR was performed to determine orientation of the MAT-containing region using primer sets a and α indicated in Fig 1C.

## Author Contributions

Conceptualization, H.M.; methodology, H.M.,N.F.; investigation, H.M., N.F.; writing, H.M.; supervision, H.M., K.T.

## Reference

Deng, Zhi-Heng, Hua-Song Ai, Cheng-Piao Lu, and Jia-Bin Li. 2020. “The Bre1/Rad6 Machinery: Writing the Central Histone Ubiquitin Mark on H2B and Beyond.” Chromosome Research: An International Journal on the Molecular, Supramolecular and Evolutionary Aspects of Chromosome Biology 28 (3–4): 247–58.

Dohmen, R. J., K. Madura, B. Bartel, and A. Varshavsky. 1991. “The N-End Rule Is Mediated by the UBC2(RAD6) Ubiquitin-Conjugating Enzyme.” Proceedings of the National Academy of Sciences of the United States of America 88 (16): 7351–55.

Feng, Dahao, Anton Stoyanov, Juliana C. Olliff, Kenneth H. Wolfe, Kantcho Lahtchev, and Sara J. Hanson. 2020. “Carbon Source Requirements for Mating and Mating-Type Switching in the Methylotrophic Yeasts Ogataea (Hansenula) Polymorpha and Komagataella Phaffii (Pichia Pastoris).” Yeast (Chichester, England) 37 (2): 237–45.

Freire-Benéitez, Verónica, R. Jordan Price, and Alessia Buscaino. 2016. “The Chromatin of Candida Albicans Pericentromeres Bears Features of Both Euchromatin and Heterochromatin.” Frontiers in Microbiology 7 (May): 759.

Guin, Krishnendu, Lakshmi Sreekumar, and Kaustuv Sanyal. 2020. “Implications of the Evolutionary Trajectory of Centromeres in the Fungal Kingdom.” Annual Review of Microbiology 74 (September): 835–53.

Hanson, Sara J., Kevin P. Byrne, and Kenneth H. Wolfe. 2014. “Mating-Type Switching by Chromosomal Inversion in Methylotrophic Yeasts Suggests an Origin for the Three-Locus Saccharomyces Cerevisiae System.” Proceedings of the National Academy of Sciences of the United States of America 111 (45): E4851–8.

Hanson, Sara J., Kevin P. Byrne, and Kenneth H. Wolfe. 2017. “Flip/Flop Mating-Type Switching in the Methylotrophic Yeast Ogataea Polymorpha Is Regulated by an Efg1-Rme1-Ste12 Pathway.” PLoS Genetics 13 (11): e1007092.

Hoege, Carsten, Boris Pfander, George-Lucian Moldovan, George Pyrowolakis, and Stefan Jentsch. 2002. “RAD6-Dependent DNA Repair Is Linked to Modification of PCNA by Ubiquitin and SUMO.” Nature 419 (6903): 135–41.

Janke, Carsten, Maria M. Magiera, Nicole Rathfelder, Christof Taxis, Simone Reber, Hiromi Maekawa, Alexandra Moreno-Borchart, et al. 2004. “A Versatile Toolbox for PCR-Based Tagging of Yeast Genes: New Fluorescent Proteins, More Markers and Promoter Substitution Cassettes.” Yeast (Chichester, England) 21 (11): 947–62.

Jentsch, S., J. P. McGrath, and A. Varshavsky. 1987. “The Yeast DNA Repair Gene RAD6 Encodes a Ubiquitin-Conjugating Enzyme.” Nature 329 (6135): 131–34.

Kao, Cheng-Fu, Cory Hillyer, Toyoko Tsukuda, Karl Henry, Shelley Berger, and Mary Ann Osley. 2004. “Rad6 Plays a Role in Transcriptional Activation through Ubiquitylation of Histone H2B.” Genes & Development 18 (2): 184–95.

Ketel, Carrie, Helen S. W. Wang, Mark McClellan, Kelly Bouchonville, Anna Selmecki, Tamar Lahav, Maryam Gerami-Nejad, and Judith Berman. 2009. “Neocentromeres Form Efficiently at Multiple Possible Loci in Candida Albicans.” PLoS Genetics 5 (3): e1000400.

Kiel, Jan A. K. W., Vladimir I. Titorenko, Ida J. van der Klei, and Marten Veenhuis. 2007. “Overproduction of Translation Elongation Factor 1-Î± (EEF1A) Suppresses the Peroxisome Biogenesis Defect in aHansenula Polymorpha Pex3Mutant via Translational Read-Through.” FEMS Yeast Research 7 (7): 1114–25.

Krassowski, Tadeusz, Jacek Kominek, Xing-Xing Shen, Dana A. Opulente, Xiaofan Zhou, Antonis Rokas, Chris Todd Hittinger, and Kenneth H. Wolfe. 2019. “Multiple Reinventions of Mating-Type Switching during Budding Yeast Evolution.” Current Biology: CB 29 (15): 2555–2562.e8.

Lee, Cheng-Sheng, and James E. Haber. 2015. “Mating-Type Gene Switching in Saccharomyces Cerevisiae.” Microbiology Spectrum 3 (2): MDNA3-0013– 2014.

Maekawa, Hiromi, Shen Jiangyan, Kaoru Takegawa, and Gislene Pereira. 2022. “SIN-like Pathway Kinases Regulate the End of Mitosis in the Methylotrophic Yeast Ogataea Polymorpha.” Cells (Basel, Switzerland) 11 (9): 1519.

Maekawa, Hiromi, and Yoshinobu Kaneko. 2014. “Inversion of the Chromosomal Region between Two Mating Type Loci Switches the Mating Type in Hansenula Polymorpha.” PLoS Genetics 10 (11): e1004796.

Obuse, Chikashi, and Jun-Ichi Nakayama. 2025. “Functional Involvement of RNAs and Intrinsically Disordered Proteins in the Assembly of Heterochromatin.” Biochimica et Biophysica Acta. General Subjects 1869 (6): 130790.

Sherman, F. 1991. “Getting Started with Yeast.” Methods in Enzymology 194: 3–21.

Sreekumar, Lakshmi, Priya Jaitly, Yao Chen, Bhagya C. Thimmappa, Amartya Sanyal, and Kaustuv Sanyal. 2019. “Cis- and Trans-Chromosomal Interactions Define Pericentric Boundaries in the Absence of Conventional Heterochromatin.” Genetics 212 (4): 1121–32.

Sun, Zu-Wen, and C. David Allis. 2002. “Ubiquitination of Histone H2B Regulates H3 Methylation and Gene Silencing in Yeast.” Nature 418 (6893): 104–8.

Sung, P., E. Berleth, C. Pickart, S. Prakash, and L. Prakash. 1991. “Yeast RAD6 Encoded Ubiquitin Conjugating Enzyme Mediates Protein Degradation Dependent on the N-End-Recognizing E3 Enzyme.” The EMBO Journal 10 (8): 2187–93.

Tanaka, Seiji, Mayumi Miyazawa-Onami, Tetsushi Iida, and Hiroyuki Araki. 2015. “IAID: An Improved Auxin-Inducible Degron System for the Construction of a ‘tight’ Conditional Mutant in the Budding Yeast Saccharomyces Cerevisiae: Improved Auxin-Inducible Degron InS. Cerevisiae.” Yeast (Chichester, England) 32 (8): 567–81.

Watkins, J. F., P. Sung, S. Prakash, and L. Prakash. 1993. “The Extremely Conserved Amino Terminus of RAD6 Ubiquitin-Conjugating Enzyme Is Essential for Amino-End Rule-Dependent Protein Degradation.” Genes & Development 7 (2): 250–61.

Wolfe, Kenneth H., and Geraldine Butler. 2022. “Mating-Type Switching in Budding Yeasts, from Flip/Flop Inversion to Cassette Mechanisms.” Microbiology and Molecular Biology Reviews: MMBR 86 (2): e0000721.

Xiao, Tiaojiang, Cheng-Fu Kao, Nevan J. Krogan, Zu-Wen Sun, Jack F. Greenblatt, Mary Ann Osley, and Brian D. Strahl. 2005. “Histone H2B Ubiquitylation Is Associated with Elongating RNA Polymerase II.” Molecular and Cellular Biology 25 (2): 637–51.

Yadav, Pawan, Manish Gupta, Rushna Wazahat, Zeyaul Islam, Susan E. Tsutakawa, Mohan Kamthan, and Pankaj Kumar. 2024. “Structural Basis for the Role of C-Terminus Acidic Tail of Saccharomyces Cerevisiae Ubiquitin-Conjugating Enzyme (Rad6) in E3 Ligase (Bre1) Mediated Recognition of Histones.” International Journal of Biological Macromolecules 254 (Pt 2): 127717.

Yamamoto, Katsuyoshi, Thi N. M. Tran, Kaoru Takegawa, Yoshinobu Kaneko, and Hiromi Maekawa. 2017. “Regulation of Mating Type Switching by the Mating Type Genes and RME1 in Ogataea Polymorpha.” Scientific Reports 7 (1): 16318.

