## Supplemental Figure 1 for "Centromere chromatin may regulate the Flip-flop type of the Mating Type Switching in methylotrophic yeast *Ogataea polymorpha*"

Supplementary Figure S1

pHM1092

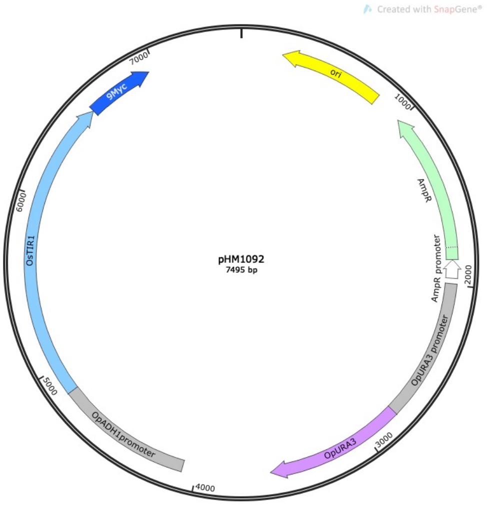


CTGCATTAATGAATCGgccaacgcgcggggagaggcggtttgcgtattgggcgctcttccgcttcctcgctcactgactcgctgcgctcggtcgttcggctgcggcgagcggtatcagctcactcaaaggcggtaatacggttatccacagaatcaggggataacgc
aggaaagaacatgtgagcaaaaggccagcaaaaggccaggaaccgtaaaaaggccgcgttgctggcgtttttccataggctcggcccccctgacgagcatcacaaaaatcgacgctcaagtcagaggtggcgaaacccgacaggactataaagataccaggcgttcc
cccctggaagctccctcgtgcgctctcctgttccgaccctgccgcttaccggatacctgtccgcctttctcccttcgggaagcgtggcgctttctcaatgctcacgctgtaggtatctcagttcggtgtaggtcgttcgctccaagctgggctgtgtgcacgaaccccccgttcagcccga
ccgctgcgccttatccggtaactatcgtcttgagtccaacccggtaagacacgacttatcgccactggcagcagccactggtaacaggattagcagagcgaggtatgtaggcggtgctacagagttcttgaagtggtggcctaactacggctacactagaaggacagtatttggt
atctgcgctctgctgaagccagttaccttcggaaaaagagttggtagctcttgatccggcaaacaaaccaccgctggtagcggtggtttttttgtttgcaagcagcagattacgcgcagaaaaaaaggatctcaagaagatcctttgatcttttctacggggtctgacgctcagtgga
acgaaaactcacgttaagggattttggtcatgagattatcaaaaaggatcttcacctagatccttttaaattaaaaatgaagttttaaatcaatctaaagtatatatgagtaaacttggtctgacagttaccaatgcttaatcagtgaggcacctatctcagcgatctgtctatttcgttcatc
catagttgcctgactgcccgtcgtgtagataactacgatacgggagggcttaccatctggccccagtgctgcaatgataccgcgagacccacgctcaccggctccagatttatcagcaataaaccagccagccggaagggccgagcgcagaagtggtcctgcaactttatcc
gcctccatccagtctattaattgttgccgggaagctagagtaagtagttcgccagttaatagtttgcgcaacgttgttgccattgctacaggcatcgtggtgtcacgctcgtcgtttggtatggcttcattcagctccggttcccaacgatcaaggcgagttacatgatcccccatgttgtga
aaaaaagcggttagctccttcggtcctccgatcgttgtcagaagtaagttggccgcagtgttatcactcatggttatggcagcactgcataattctcttactgtcatgccatccgtaagatgcttttctgtgactggtgagtactcaaccaagtcattctgagaatagtgtatgcggcgacc
gagttgctcttgcccggcgtcaatacgggataataccgcgccacatagcagaactttaaaagtgctcatcattggaaaacgttcttcggggcgaaaactctcaaggatcttaccgctgttgagatccagttcgatgtaacccactcgtgcacccaactgatcttcagcatcttttacttt
caccagcgtttctgggtgagcaaaaacaggaaggcaaaatgccgcaaaaaagggaataagggcgacacggaaatgttgaatactcatactcttcctttttcaatattattgaagcatttatcagggttattgtctcatgagcggatacatatttgaatgtatttagaaaaataaaca
aataggggttccgcgcacatttccccgaaaagtgccacctgacgtcTCCAGAACCTGCTACACCATCTACGCCAGCAAGTGGAGATGAAGAACGTGGGAGCAAGCGACGCGGATCCGAAAGCGTCCCCCCTGG
TTGTACTAAGCGCTTCAATGCTCTGTCTGCACAACTTCTCACACAGATATCATCCAACCGGTTTGCGTCTATGTTCATGCAACCAGTGAATGAAAACGAAGAGCCCAACTACTACAAGCTAAT
ACGGCATCCCGTGGACATTAAGAGTTTGGGTAAAGCTATTAGGACGGGTGAAATAAAGTCATTCGACGAGTTGGAATTTGAGCTTCAACTGATGTTCAGCAACGCAATTATGTACAACGATA
TGAATCAGACAGAAACTTACAAATGGACGATTGAAATGATGGAAGAAACGCAAAACTTGTTGAGCCTATTCCGGGAATCATCCAGCAATTAAAGCTCATTAATTGCCGCCTCTCATCTGGTTC
ATCAACATCTGGTTCTGTTCCTGGAATCTTAAACCAACTCTTTCGGAGTGCTTGAGGAACTTTTCAGCGCACTTGTTCAAACATGTTTCTTCTCTAGAGCTAAGTTTGTTGGAGGTGAAGTCG
TTGACGCAATCATTAAAACATCTGTCCACCAAACTGGTGTAGAGCTGCAATGTTAGTTTTCAGAAGGCGTTCACTTACTCTCATGAAGTCATTCATTTGCTTCTGCTCCACAATTTTCTGGAAT
TCTTGCTGTTCTTTATAGTTGAGTTGATCCATGGTGTATCAAAAATAAACTAATGAATTATCGAAAAATTTTCAACATTTCCCTGAATAATAATCAACATGTATAAATCTTATGGAGAAAGGGCG
AAGTCTCACCCATCTAAGGTCGCCAGCAGACTACTTAATTTGATGGAATCCAAGCAAACAAACCTCTGCGCTTCTGTGGATGTGACTAAAACTCAGGAATTATTGGAGCTTCTTGATAAACTG
GGCCCTTACATCTGCCTTGTCAAAACTCATATTGACATAGTAGAGGACTTCTCTTATGAACACACCATTTTACCATTACAAGGACTTGCAAAGAAACACAACTTCATGATTTTTGAAGACAGAA
AGTTTGCTGATATAGGAAACACAGTCAAACTACAGTATAAGGGAGGAATTTATCGAACATCCAAGTGGGCCGATATCACGAATGCACACGGAGTGACTGGTGCAGGAATTGTTGAAGGTCTT
AAACAGGCCGCAGAAGAAAGTACAGATGAGCCACGTGGGCTTTTGATGCTTGCTGAGCTCTCTTCAAAGGGATCATTAGCTACCGGTGAGTATACTCAAAAAACTGTGGAAATAGCGAAAA
GCGATAAAGAATTTGTCATTGGATTTATTGCACAGAGAGACATGGGAGGTCGTGAGGAAGGCTTTGACTGGCTGATCATGACTCCAGGAGTTGGTTTAGATGATAAAGGCGATTCTCTGGG
CCAACAGTACAGAACTGTTGATGAAGTGATGCAAACAGGAACCGATGTCATTATCGTTGGAAGAGGTTTATTCGGAAAAGGAAGAGATCCTGAAGTGGAAGGGAAGAGATACAGAAATGCT
GGGTGGGAAGCTTACAAGCGGCGCATTGCTTAACGGCTTTCAGTTCTATATACATCGTCAAAATTGATTTTCGCTAAAATGCTGACGGGATATTTCGAATACGAAAAGCCCAATAGAAGTCG
CGGGAATACTAGTGAAGGACGATCTGAAAAAGCCTCTATAGCAGAAGCCAAAGAGGGAGAGGTGAGATTTTCGCCATCGTTGTAGAATAATATCAAATGTCTCAATGTAAGAATTATAGTAG
AGTTTGAAGAAGCGACGCCGATCCAGTTGATGTGCTTCGTTGTAGAGATCTAGAGATTCGAGTCTAGTCAGGTGGACCTTTTGAATTTTATTGAGTGGATATTGGATTGCAAGCAGTGTAGTT
GCGGCGGAAGCTCCAGCAAGAAGGATAAACGTCAAATTCAAAGCTTTGAAAAATCTTGAATTCTTCTCCTGGTTTTCTTCATCATGAAAAAGTTTCGAGCGCCAATATTGATACCACTGTATA
GCTGCTTTTGTCTTGTGGTACCCCTGGTTTTTCACAAGTTCATTTGTCACACGGCACCTCGAAGTTGCACTTAACTTTGCGCTTCTTGCAGCTGAGACATGTCTTGGAAATACGAGTAGTACG
CCGTAGCTTGGTTATGTCGAGCCTAGTAAACTCCGCAGGAGAATTCCTCTTCTCCGAAGGCACTCTCTCCGCAGAAGCCATAAATTATTTGAAATCAGAAATGTGAGCGATATAAACACCCC
TGCACGGGGCTGCACCCCGATTTTTTTCGGGTGATCGTATGATGCTACTATGAGCCCGCAGATAACATCTGACTTTACATTAAGGCAAAATTCTGGTGTAGGGTGTCCGCAGGCCGAGTAA
TGCTGACCGGTACGCACCTTTTTCGTCCGTTGTGGTGTGCTGCTTCATGGTCAATTTTTTTTAGCTCAGATGACATCGGAAGGTGTTCCTACAAAGCACACCTCCAATTCGCCGTAAAACCT
CGGAGTAAATCATTCTCCCTGTAGTGTAAACTAAGCTGATGAGGCGTTCGGCAGATCCAATTGGACACACCCCGTCATGGGGTGTGAAATACCCCGCCACGGCCATCCGGCAGCCCCACT
CCACCACTTTGGGTTGAATATGGTCCAATTGTTATGTGCAAATTTTCGGACATTGGAAATCACCCACTCGGAGTAAAGCAACTCGTAGTGACTGAAAAATAAGACGTCATCATTAATTAGGCT
GCAACAGACATATAAATACGAGACACTTTTCCCTCTTCTTAGGCATCCTTCTATCAATCAATCAATCAATTTAAAATGacgtacttcccggaggaGGTGGTGGAGCACATCTTCAGCTTCCTGCCGGC
GCAGCGCGACCGCAACACGGTCTCGCTCGTCTGCAAGGTGTGGTACGAGATCGAGAGGCTGAGCCGCCGCGGCGTCTTCGTGGGCAACTGCTACGCCGTGCGCGCCGGCCGCGTCGC
CGCGCGGTTCCCCAACGTGCGGGCGCTCACGGTGAAGGGGAAGCCCCACTTCGCCGACTTCAACCTCGTGCCCCCCGACTGGGGCGGCTACGCGGGGCCGTGGATCGAGGCGGCCG
CGAGGGGATGCCACGGCCTGGAGGAGCTCAGGATGAAGCGGATGGTGGTGTCCGACGAGAGCCTCGAGCTGCTGGCTCGCTCGTTCCCGCGGTTCAGGGCTCTTGTTCTTATCAGCTG
CGAGGGGTTCAGCACTGACGGGCTAGCCGCCGTCGCGAGCCATTGCAAGCTTCTGAGGGAGTTGGATTTGCAGGAAAATGAAGTGGAGGATCGAGGGCCTAGGTGGCTTTCCTGCTTCC
CTGATTCCTGCACATCACTTGTCTCATTGAATTTTGCCTGCATCAAAGGGGAGGTTAATGCTGGTTCACTGGAGAGACTTGTTAGCAGGTCCCCAAACCTGCGGAGTTTGAGGCTGAATCGA
TCTGTATCGGTAGATACACTTGCAAAGATACTACTGCGTACCCCTAACTTGGAGGATTTGGGGACAGGGAATTTGACAGATGACTTCCAAACTGAGTCCTACTTTAAGCTTACCAGTGCTCT
GGAGAAATGCAAGATGTTGAGGAGTTTGTCTGGATTCTGGGATGCTTCTCCTGTTTGCCTGTCATTTATCTACCCCCTGTGTGCTCAACTGACAGGATTGAACTTGAGCTATGCCCCCACAC
TTGATGCTTCTGACCTTACAAAAATGATTAGCCGCTGTGTGAAGCTCCAACGCCTTTGGGTACTGGATTGTATCTCGGACAAAGGCTTGCAAGTGGTGGCCTCCAGTTGCAAAGACTTGCAA
GAACTCAGGGTATTTCCATCAGATTTCTACGTAGCTGGTTATTCTGCAGTGACAGAGGAGGGACTTGTTGCAGTATCCTTGGGCTGTCCAAAACTGAACTCACTACTGTACTTCTGTCACCA
AATGACTAATGCTGCACTAGTTACTGTCGCCAAGAACTGTCCAAATTTCACACGATTCAGACTTTGTATTCTTGAGCCAGGGAAGCCTGATGTTGTGACAAGCCAACCATTAGATGAAGGCT
TTGGAGCTATTGTTCGTGAGTGCAAGGGATTACAACGTTTGTCAATATCTGGTCTTCTCACAGACAAAGTTTTCATGTATATTGGGAAATATGCAAAACAACTTGAGATGCTTTCTATAGCATT
TGCTGGTGACAGTGATAAGGGTATGATGCATGTTATGAATGGATGCAAGAATTTAAGGAAACTGGAGATAAGAGATAGCCCGTTTGGTGATGCTGCACTCTTGGGGAATTTTGCTAGGTACG
AGACAATGCGATCCCTTTGGATGTCATCTTGCAATGTCACGTTAAAGGGGTGCCAAGTCCTTGCGTCAAAGATGCCGATGCTCAATGTTGAGGTCATAAATGAGCGGGATGGTAGCAATGA
AATGGAGGAAAACCATGGAGATCTGCCTAAAGTGGAGAAATTATATGTGTACCGCACAACTGCTGGGGCGAGGGATGATGCACCAAATTTTGTTAAAATCCTAgtcgacTCCGGTTCTGCTGC
TAGTGGTGAACAAAAGTTGATTTCTGAAGAAGATTTGAACGGGGAACAAAAGCTAATCTCCGAGGAAGACTTGAACGGTGAACAAAAATTAATCTCAGAAGAAGACTTGAACGGATCCTCTA
GAGGTGAACAAAAGTTGATTTCTGAAGAAGATTTGAACGGTGAACAAAAGCTAATCTCCGAGGAAGACTTGAACGGTGAACAAAAATTAATCTCAGAAGAAGACTTGAACGGATCCTCTAGA
GGTGAACAAAAGTTGATTTCTGAAGAAGATTTGAACGGTGAACAAAAGCTAATCTCCGAGGAAGACTTGAACGGTGAACAAAAATTAATCTCAGAAGAAGACTTGAACGGATCCACTAGCTA
AAAAGTggaacgatcattcaAGAGAtCCGGGccgataccctaatgagatcttccgttatataaccatcgataacctgaataaaaaatctgctgtcaaagccgtctgcctggcaccagggcgaatatgatgctcagaacagctgtcatcagcaatatgagcaacagca
caaagatggagcggcgcttcagcaaattcttcacgaacggctccttgatgcacacaagcccattcttcgatgcgtagctctcgctgtagcccgcaaaatatagcaaggggtctacccggtagtagctgcttagcagcaagtagaatgcaaaaaagaacggcacaacaacga
cccagcgcatctgctttggttgatagtcaaggaaaataagtgtatatccggtccagaaagcccccaatccagaaaaaaacgagcccaacattctataggctgctgattttgacgttacgtttgacaatgcatgatatcggaggaaattagggtatgcctcgtgttccatagccttaa
aaacaaactcccgcgaCTCGAG

### pHM1217

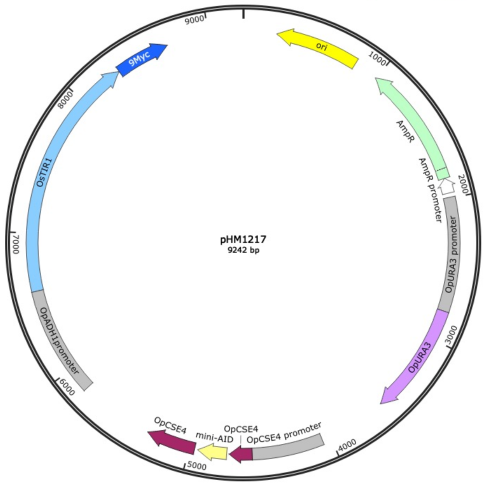


CTGCATTAATGAATCGgccaacgcgcggggagaggcggtttgcgtattgggcgctcttccgcttcctcgctcactgactcgctgcgctcggtcgttcggctgcggcgagcggtatcagctcactcaaaggcggtaatacggttatccacagaatca
ggggataacgcaggaaagaacatgtgagcaaaaggccagcaaaaggccaggaaccgtaaaaaggccgcgttgctggcgtttttccataggctcggcccccctgacgagcatcacaaaaatcgacgctcaagtcagaggtggcgaaacccgaca
ggactataaagataccaggcgttcccccctggaagctccctcgtgcgctctcctgttccgaccctgccgcttaccggatacctgtccgcctttctcccttcgggaagcgtggcgctttctcaatgctcacgctgtaggtatctcagttcggtgtaggtcgttcgctcc
aagctgggctgtgtgcacgaaccccccgttcagcccgaccgctgcgccttatccggtaactatcgtcttgagtccaacccggtaagacacgacttatcgccactggcagcagccactggtaacaggattagcagagcgaggtatgtaggcggtgctacag
agttcttgaagtggtggcctaactacggctacactagaaggacagtatttggtatctgcgctctgctgaagccagttaccttcggaaaaagagttggtagctcttgatccggcaaacaaaccaccgctggtagcggtggtttttttgtttgcaagcagcagattac
gcgcagaaaaaaaggatctcaagaagatcctttgatcttttctacggggtctgacgctcagtggaacgaaaactcacgttaagggattttggtcatgagattatcaaaaaggatcttcacctagatccttttaaattaaaaatgaagttttaaatcaatctaaagt
atatatgagtaaacttggtctgacagttaccaatgcttaatcagtgaggcacctatctcagcgatctgtctatttcgttcatccatagttgcctgactgcccgtcgtgtagataactacgatacgggagggcttaccatctggccccagtgctgcaatgataccgcg
agacccacgctcaccggctccagatttatcagcaataaaccagccagccggaagggccgagcgcagaagtggtcctgcaactttatccgcctccatccagtctattaattgttgccgggaagctagagtaagtagttcgccagttaatagtttgcgcaacgt
tgttgccattgctacaggcatcgtggtgtcacgctcgtcgtttggtatggcttcattcagctccggttcccaacgatcaaggcgagttacatgatcccccatgttgtgaaaaaaagcggttagctccttcggtcctccgatcgttgtcagaagtaagttggccgcagt
gttatcactcatggttatggcagcactgcataattctcttactgtcatgccatccgtaagatgcttttctgtgactggtgagtactcaaccaagtcattctgagaatagtgtatgcggcgaccgagttgctcttgcccggcgtcaatacgggataataccgcgccaca
tagcagaactttaaaagtgctcatcattggaaaacgttcttcggggcgaaaactctcaaggatcttaccgctgttgagatccagttcgatgtaacccactcgtgcacccaactgatcttcagcatcttttactttcaccagcgtttctgggtgagcaaaaacagga
aggcaaaatgccgcaaaaaagggaataagggcgacacggaaatgttgaatactcatactcttcctttttcaatattattgaagcatttatcagggttattgtctcatgagcggatacatatttgaatgtatttagaaaaataaacaaataggggttccgcgcaca
tttccccgaaaagtgccacctgacgtcTCCAGAACCTGCTACACCATCTACGCCAGCAAGTGGAGATGAAGAACGTGGGAGCAAGCGACGCGGATCCGAAAGCGTCCCCCCTGGTTGTA
CTAAGCGCTTCAATGCTCTGTCTGCACAACTTCTCACACAGATATCATCCAACCGGTTTGCGTCTATGTTCATGCAACCAGTGAATGAAAACGAAGAGCCCAACTACTACAAG
CTAATACGGCATCCCGTGGACATTAAGAGTTTGGGTAAAGCTATTAGGACGGGTGAAATAAAGTCATTCGACGAGTTGGAATTTGAGCTTCAACTGATGTTCAGCAACGCAAT
TATGTACAACGATATGAATCAGACAGAAACTTACAAATGGACGATTGAAATGATGGAAGAAACGCAAAACTTGTTGAGCCTATTCCGGGAATCATCCAGCAATTAAAGCTCATT
AATTGCCGCCTCTCATCTGGTTCATCAACATCTGGTTCTGTTCCTGGAATCTTAAACCAACTCTTTCGGAGTGCTTGAGGAACTTTTCAGCGCACTTGTTCAAACATGTTTCTT
CTCTAGAGCTAAGTTTGTTGGAGGTGAAGTCGTTGACGCAATCATTAAAACATCTGTCCACCAAACTGGTGTAGAGCTGCAATGTTAGTTTTCAGAAGGCGTTCACTTACTCTC
ATGAAGTCATTCATTTGCTTCTGCTCCACAATTTTCTGGAATTCTTGCTGTTCTTTATAGTTGAGTTGATCCATGGTGTATCAAAAATAAACTAATGAATTATCGAAAAATTTTCA
ACATTTCCCTGAATAATAATCAACATGTATAAATCTTATGGAGAAAGGGCGAAGTCTCACCCATCTAAGGTCGCCAGCAGACTACTTAATTTGATGGAATCCAAGCAAACAAAC
CTCTGCGCTTCTGTGGATGTGACTAAAACTCAGGAATTATTGGAGCTTCTTGATAAACTGGGCCCTTACATCTGCCTTGTCAAAACTCATATTGACATAGTAGAGGACTTCTCT
TATGAACACACCATTTTACCATTACAAGGACTTGCAAAGAAACACAACTTCATGATTTTTGAAGACAGAAAGTTTGCTGATATAGGAAACACAGTCAAACTACAGTATAAGGGA
GGAATTTATCGAACATCCAAGTGGGCCGATATCACGAATGCACACGGAGTGACTGGTGCAGGAATTGTTGAAGGTCTTAAACAGGCCGCAGAAGAAAGTACAGATGAGCCA
CGTGGGCTTTTGATGCTTGCTGAGCTCTCTTCAAAGGGATCATTAGCTACCGGTGAGTATACTCAAAAAACTGTGGAAATAGCGAAAAGCGATAAAGAATTTGTCATTGGATTT
ATTGCACAGAGAGACATGGAGGTCGTGAGGAAGGCTTTGACTGGCTGATCATGACTCCAGGAGTTGGTTTAGATGATAAAGGCGATTCTCTGGGCCAACAGTACAGAACT
GTTGATGAAGTGATGCAAACAGGAACCGATGTCATTATCGTTGGAAGAGGTTTATTCGGAAAAGGAAGAGATCCTGAAGTGGAAGGGAAGAGATACAGAAATGCTGGGTGGG
AAGCTTACAAGCGGCGCATTGCTTAACGGCTTTCAGTTCTATATACATCGTCAAAATTGATTTTCGCTAAAATGCTGACGGGATATTTCGAATACGAAAAGCCCAATAGAAGTC
GCGGGAATACTAGTGAAGGACGATCTGAAAAAGCCTCTATAGCAGAAGCCAAAGAGGGAGAGGTGAGATTTTCGCCATCGTTGTAGAATAATATCAAATGTCTCAATGTAAGA
ATTATAGTAGAGTTTGAAGAAGCGACGCCGATCCAGTTGATGTGCTTCGTTGTAGAGATCTAGAGATTCGAGTCTAGTCAGGTGGACCTTTTGAATTTTATTGAGTGGATATTG
GATTGCAAGCAGTGTAGTTGCGGCGGAAGCTCCAGCAAGAAGGATAAACGTCAAATTCAAAGCTTTGAAAAATCTTGAATTCTTCTCCTGGTTTTCTTCATCATGAAAAAGTTT
CGAGCGCCAATATTGATACCACTGTATAGCTGCTTTTGTCTTGTGGACGAGCCGAAGCTGGTGAAGCCTAAGAAGGAACAGCAAGAGGAGCCGGAGGAGCCAGAGGAGCC
AACGGCAGATGTGGCTGCCAGCCAGCCAGCTGCAGTCGAGAAACATGCTGCTGCGGACGAAAAGCCATCTAAGAAGGGCAAGGAGGGTCTGGGAAAACGGTTCCTCAAGT
TTATGACCCAGGAACCACCAAAGCCAAAGAAGAAGGTGAAGGGCTCTCCCAAAACAGCTTCTAAGACCGCTCCTAAACCGAAGGAGACCAAGAAAGAGACCACAGAGGAGG
CGGGAGATGCATTCTCAAACTTCTCCCAGGGCTCAGAGGTCGCCGAGAAATAGGACTGTTCATTTCCTGTGCATTTATTTGTATTATGTAGGTGAAAACATATTGAAGAAATTT
ATTAATTTTTTAAATATTTTTTGAGACCAGCCACTGCACTTAATTATTTCCAAAGGGGTAATCAAAAAATTGTCCTGATAAACTTTAGGGCATTGTTTCAATGGCCAGGTTAAATA
AGCACAAGCCGGTGTCGACAACGCCAAAGACGCCCAAGATGGGCTCCAGCACAAGCTTCAAGAATTCGCCAGCCAGTATCAAGAAAACACCGTCTAAAAGCTCTGAGACGC
CAAAGGTCAAGGCGGGCTCGACGTTGAGCTCTggtgctggagcaggcgccAAAGAAAAATCTGCCTGTCCAAAAGATCCTGCAAAGCCACCTGCTAAAGCACAAGTCGTGGGTTGG
CCTCCTGTTAGAAGTTATAGAAAGAATGTGATGGTTTCTTGTCAAAAATCTAGTGGAGGTCCTGAAGCCGCAGCCTTCGTCAAAGTATCTATGGACGGAGCTCCTTATTTGAG
AAAGATAGACTTGAGAATGTATAAAggtgccggagctggagcaggtgctGGGCTCGCGAGAAACCAGCCAGGCGACCCGACAGAAATCAAAACGCAGAAAAAAAGATTCAAGCCCGGA
ACTGTGGCCTTGCGCGAGATCAGAAGGTTCCAGAAGTCCACAGAGCTGCTTATCAGGAAACTACCCTTTGCTAGACTCGTCAGAGAAATCGTCCAGGACGAGTTCGGAACA
AGCACCTACAGATGGCAGTCGGTCGCAGTGCTGGCGTTGCAGGAAGCCGTCGAGGCGTATCTTGTGCATCTTTTTGAGGATACCAACCTGTGCGCGCTCCACGCGAAGCG
AGTCACCATCATGCAGAAAGACATCCATCTTGCCAGGAGACTGCGGGGGGATTGAGTCCTATTAGTGTATGTATACTACAACCTGTTGTGTCAAATACCAATCGGGCCTTTTT
CCATGTAGTCCTCGCGCAGGAAGGAGCCCCCGACAGCGTGCGAGTTGAACACCTGCTTCGCGAGTATCTGTGCGCCCAGAAACGAGCAGTCCTCGAACCCGGTCTTTTTC
CACTCGGTAAAGTAGTCCGGCCGACTCAGTAGCTTGACGTGTTTCGGCACCTGGATTAGGTTCAGGTCCTGAATCATGTTGTTTGTGTCGCGGAACGCTCCGCTTTGGGCAG
ACATTGCACCCTGTCCGACAACAAAATTGCGGTACAGGTGCACCACGAGCGCCTCCTTGAATCCGATGATTTTGCTCGTCTGGCCTACTAGGATCAAGTTGTCGTAGCATTC
CTGCGCCCTTTTAAGATCTGTGATTTTCGACAATGTGCGGTGGATAGCCGCTCCAATGCTCTCGATCAGGTCTGTGCAGCCTTGGAACCGCTCCTTACCCACGCTCACACCC
TGGTTTTTCACAAGTTCATTTGTCACACGGCACCTCGAAGTTGCACTTAACTTTGCGCTTCTTGCAGCTGAGACATGTCTTGGAAATACGAGTAGTACGCCGTAGCTTGGTTAT
GTCGAGCCTAGTAAACTCCGCAGGAGAATTCCTCTTCTCCGAAGGCACTCTCTCCGCAGAAGCCATAAATTATTTGAAATCAGAAATGTGAGCGATATAAACACCCCTGCAC
GGGGCTGCACCCCGATTTTTTTCGGGTGATCGTATGATGCTACTATGAGCCCGCAGATAACATCTGACTTTACATTAAGGCAAAATTCTGGTGTAGGGTGTCCGCAGGCCGA
GTAATGCTGACCGGTACGCACCTTTTTCGTCCGTTGTGGTGTGCTGCTTCATGGTCAATTTTTTTTAGCTCAGATGACATCGGAAGGTGTTCCTACAAAGCACACCTCCAATT
CGCCGTAAAACCTCGGAGTAAATCATTCTCCCTGTAGTGTAAACTAAGCTGATGAGGCGTTCGGCAGATCCAATTGGACACACCCCGTCATGGGGTGTGAAATACCCCGCCA
CGGCCATCCGGCAGCCCCACTCCACCACTTTGGGTTGAATATGGTCCAATTGTTATGTGCAAATTTTCGGACATTGGAAATCACCCACTCGGAGTAAAGCAACTCGTAGTGA
CTGAAAAATAAGACGTCATCATTAATTAGGCTGCAACAGACATATAAATACGAGACACTTTTCCCTCTTCTTAGGCATCCTTCTATCAATCAATCAATCAATTTAAAATGacgtactt
cccggaggaGGTGGTGGAGCACATCTTCAGCTTCCTGCCGGCGCAGCGCGACCGCAACACGGTCTCGCTCGTCTGCAAGGTGTGGTACGAGATCGAGAGGCTGAGCCGCCG
CGGCGTCTTCGTGGGCAACTGCTACGCCGTGCGCGCCGGCCGCGTCGCCGCGCGGTTCCCCAACGTGCGGGCGCTCACGGTGAAGGGGAAGCCCCACTTCGCCGACTT
CAACCTCGTGCCCCCCGACTGGGGCGGCTACGCGGGGCCGTGGATCGAGGCGGCCGCGAGGGGATGCCACGGCCTGGAGGAGCTCAGGATGAAGCGGATGGTGGTGTC
CGACGAGAGCCTCGAGCTGCTGGCTCGCTCGTTCCCGCGGTTCAGGGCTCTTGTTCTTATCAGCTGCGAGGGGTTCAGCACTGACGGGCTAGCCGCCGTCGCGAGCCATT
GCAAGCTTCTGAGGGAGTTGGATTTGCAGGAAAATGAAGTGGAGGATCGAGGGCCTAGGTGGCTTTCCTGCTTCCCTGATTCCTGCACATCACTTGTCTCATTGAATTTTGC
CTGCATCAAAGGGGAGGTTAATGCTGGTTCACTGGAGAGACTTGTTAGCAGGTCCCCAAACCTGCGGAGTTTGAGGCTGAATCGATCTGTATCGGTAGATACACTTGCAAAG
ATACTACTGCGTACCCCTAACTTGGAGGATTTGGGGACAGGGAATTTGACAGATGACTTCCAAACTGAGTCCTACTTTAAGCTTACCAGTGCTCTGGAGAAATGCAAGATGTT
GAGGAGTTTGTCTGGATTCTGGGATGCTTCTCCTGTTTGCCTGTCATTTATCTACCCCCTGTGTGCTCAACTGACAGGATTGAACTTGAGCTATGCCCCCACACTTGATGCTT
CTGACCTTACAAAAATGATTAGCCGCTGTGTGAAGCTCCAACGCCTTTGGGTACTGGATTGTATCTCGGACAAAGGCTTGCAAGTGGTGGCCTCCAGTTGCAAAGACTTGCA
AGAACTCAGGGTATTTCCATCAGATTTCTACGTAGCTGGTTATTCTGCAGTGACAGAGGAGGGACTTGTTGCAGTATCCTTGGGCTGTCCAAAACTGAACTCACTACTGTACT
TCTGTCACCAAATGACTAATGCTGCACTAGTTACTGTCGCCAAGAACTGTCCAAATTTCACACGATTCAGACTTTGTATTCTTGAGCCAGGGAAGCCTGATGTTGTGACAAGC
CAACCATTAGATGAAGGCTTTGGAGCTATTGTTCGTGAGTGCAAGGGATTACAACGTTTGTCAATATCTGGTCTTCTCACAGACAAAGTTTTCATGTATATTGGGAAATATGCA
AAACAACTTGAGATGCTTTCTATAGCATTTGCTGGTGACAGTGATAAGGGTATGATGCATGTTATGAATGGATGCAAGAATTTAAGGAAACTGGAGATAAGAGATAGCCCGTTT
GGTGATGCTGCACTCTTGGGGAATTTTGCTAGGTACGAGACAATGCGATCCCTTTGGATGTCATCTTGCAATGTCACGTTAAAGGGGTGCCAAGTCCTTGCGTCAAAGATGC
CGATGCTCAATGTTGAGGTCATAAATGAGCGGGATGGTAGCAATGAAATGGAGGAAAACCATGGAGATCTGCCTAAAGTGGAGAAATTATATGTGTACCGCACAACTGCTGG
GGCGAGGGATGATGCACCAAATTTTGTTAAAATCCTAgtcgacTCCGGTTCTGCTGCTAGTGGTGAACAAAAGTTGATTTCTGAAGAAGATTTGAACGGGGAACAAAAGCTAATC
TCCGAGGAAGACTTGAACGGTGAACAAAAATTAATCTCAGAAGAAGACTTGAACGGATCCTCTAGAGGTGAACAAAAGTTGATTTCTGAAGAAGATTTGAACGGTGAACAAAA
GCTAATCTCCGAGGAAGACTTGAACGGTGAACAAAAATTAATCTCAGAAGAAGACTTGAACGGATCCTCTAGAGGTGAACAAAAGTTGATTTCTGAAGAAGATTTGAACGGTG
AACAAAAGCTAATCTCCGAGGAAGACTTGAACGGTGAACAAAAATTAATCTCAGAAGAAGACTTGAACGGATCCACTAGCTAAAAAGTggaacgatcattcaAGAGAtCCGGGccgata
ccctaatgagatcttccgttatataaccatcgataacctgaataaaaaatctgctgtcaaagccgtctgcctggcaccagggcgaatatgatgctcagaacagctgtcatcagcaatatgagcaacagcacaaagatggagcggcgcttcagcaaattctt
cacgaacggctccttgatgcacacaagcccattcttcgatgcgtagctctcgctgtagcccgcaaaatatagcaaggggtctacccggtagtagctgcttagcagcaagtagaatgcaaaaaagaacggcacaacaacgacccagcgcatctgctttgg
ttgatagtcaaggaaaataagtgtatatccggtccagaaagcccccaatccagaaaaaaacgagcccaacattctataggctgctgattttgacgttacgtttgacaatgcatgatatcggaggaaattagggtatgcctcgtgttccatagccttaaaaaca
aactcccgcgaCTCGAG

### pHM1219

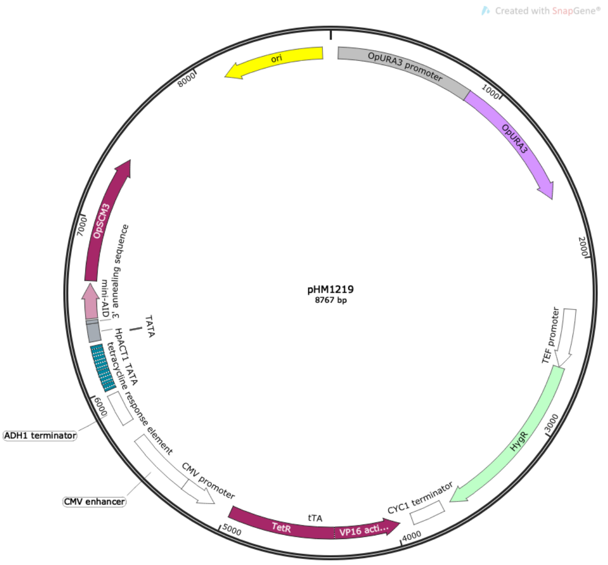


ACGCGGCCGCCAGCTGAAGCTTCGTACGCTGCAGGTCGACGGATCTCCAGAACCTGCTACACCATCTACGCCAGCAAGTGGAGATGAAGAACGTGGGAGCAAGCGACGCGGATCCGA
AAGCGTCCCCCCTGGTTGTACTAAGCGCTTCAATGCTCTGTCTGCACAACTTCTCACACAGATATCATCCAACCGGTTTGCGTCTATGTTCATGCAACCAGTGAATGAAAACGAAGAGCCCA
ACTACTACAAGCTAATACGGCATCCCGTGGACATTAAGAGTTTGGGTAAAGCTATTAGGACGGGTGAAATAAAGTCATTCGACGAGTTGGAATTTGAGCTTCAACTGATGTTCAGCAACGCAAT
TATGTACAACGATATGAATCAGACAGAAACTTACAAATGGACGATTGAAATGATGGAAGAAACGCAAAACTTGTTGAGCCTATTCCGGGAATCATCCAGCAATTAAAGCTCATTAATTGCCGCC
TCTCATCTGGTTCATCAACATCTGGTTCTGTTCCTGGAATCTTAAACCAACTCTTTCGGAGTGCTTGAGGAACTTTTCAGCGCACTTGTTCAAACATGTTTCTTCTCTAGAGCTAAGTTTGTTGGA
GGTGAAGTCGTTGACGCAATCATTAAAACATCTGTCCACCAAACTGGTGTAGAGCTGCAATGTTAGTTTTCAGAAGGCGTTCACTTACTCTCATGAAGTCATTCATTTGCTTCTGCTCCACAATT
TTCTGGAATTCTTGCTGTTCTTTATAGTTGAGTTGATCCATGGTGTATCAAAAATAAACTAATGAATTATCGAAAAATTTTCAACATTTCCCTGAATAATAATCAACATGTATAAATCTTATGGAGAAAG
GGCGAAGTCTCACCCATCTAAGGTCGCCAGCAGACTACTTAATTTGATGGAATCCAAGCAAACAAACCTCTGCGCTTCTGTGGATGTGACTAAAACTCAGGAATTATTGGAGCTTCTTGATAA
ACTGGGCCCTTACATCTGCCTTGTCAAAACTCATATTGACATAGTAGAGGACTTCTCTTATGAACACACCATTTTACCATTACAAGGACTTGCAAAGAAACACAACTTCATGATTTTTGAAGACA
GAAAGTTTGCTGATATAGGAAACACAGTCAAACTACAGTATAAGGGAGGAATTTATCGAACATCCAAGTGGGCCGATATCACGAATGCACACGGAGTGACTGGTGCAGGAATTGTTGAAGGT
CTTAAACAGGCCGCAGAAGAAAGTACAGATGAGCCACGTGGGCTTTTGATGCTTGCTGAGCTCTCTTCAAAGGGATCATTAGCTACCGGTGAGTATACTCAAAAAACTGTGGAAATAGCGAA
AAGCGATAAAGAATTTGTCATTGGATTTATTGCACAGAGAGACATGGAGGTCGTGAGGAAGGCTTTGACTGGCTGATCATGACTCCAGGAGTTGGTTTAGATGATAAAGGCGATTCTCTGGG
CCAACAGTACAGAACTGTTGATGAAGTGATGCAAACAGGAACCGATGTCATTATCGTTGGAAGAGGTTTATTCGGAAAAGGAAGAGATCCTGAAGTGGAAGGGAAGAGATACAGAAATGCTG
GGTGGGAAGCTTACAAGCGGCGCATTGCTTAACGGCTTTCAGTTCTATATACATCGTCAAAATTGATTTTCGCTAAAATGCTGACGGGATATTTCGAATACGAAAAGCCCAATAGAAGTCGCG
GGAATACTAGTGAAGGACGATCTGAAAAAGCCTCTATAGCAGAAGCCAAAGAGGGAGAGGTGAGATTTTCGCCATCGTTGTAGAATAATATCAAATGTCTCAATGTAAGAATTATAGTAGAGTT
TGAAGAAGCGACGCCGATCCAGTTGATGTGCTTCGTTGTAGAGATCTAGAGATTCGAGTCTAGTCAGGTGGACCTTTTGAATTTTATTGAGTGGATATTGGATTGCAAGCAGTGTAGTTGCGGC
GGAAGCTCCAGCAAGAAGGATAAACGTCAAATTCAAAGCTTTGAAAAATCTTGAATTCTTCTCCTGGTTTTCTTCATCATGAAAAAGTTTCGAGCGCCAATATTGATACCACTGTATAGCTGCTTT
TGTCTTGTGGTACCCCTGGTTTTTCACAAGTTCggcccaaacttgatggatctttggaggctcgaacagttcttgaggttaccctaacaagttaagaacgattattatatgcagccgttaGGGTTAATTAAGGCGCGCCAGATC
TGTTTAGCTTGCCTCGTCCCCGCCGGGTCACCCGGCCAGCGACATGGAGGCCCAGAATACCCTCCTTGACAGTCTTGACGTGCGCAGCTCAGGGGCATGATGTGACTGTCGCCCGTAC
ATTTAGCCCATACATCCCCATGTATAATCATTTGCATCCATACATTTTGATGGCCGCACGGCGCGAAGCAAAAATTACGGCTCCTCGCTGCAGACCTGCGAGCAGGGAAACGCTCCCCTCA
CAGACGCGTTGAATTGTCCCCACGCCGCGCCCCTGTAGAGAAATATAAAAGGTTAGGATTTGCCACTGAGGTTCTTCTTTCATATACTTCCTTTTAAAATCTTGCTAGGATACAGTTCTCACATC
ACATCCGAACATAAACAACCATGGGTAAAAAGCCTGAACTCACCGCGACGTCTGTCGAGAAGTTTCTGATCGAAAAGTTCGACAGCGTCTCCGACCTGATGCAGCTCTCGGAGGGCGAAG
AATCTCGTGCTTTCAGCTTCGATGTAGGAGGGCGTGGATATGTCCTGCGGGTAAATAGCTGCGCCGATGGTTTCTACAAAGATCGTTATGTTTATCGGCACTTTGCATCGGCCGCGCTCCCG
ATTCCGGAAGTGCTTGACATTGGGGAATTCAGCGAGAGCCTGACCTATTGCATCTCCCGCCGTGCACAGGGTGTCACGTTGCAAGACCTGCCTGAAACCGAACTGCCCGCTGTTCTGCAG
CCGGTCGCGGAGGCCATGGATGCGATCGCTGCGGCCGATCTTAGCCAGACGAGCGGGTTCGGCCCATTCGGACCGCAAGGAATCGGTCAATACACTACATGGCGTGATTTCATATGCG
CGATTGCTGATCCCCATGTGTATCACTGGCAAACTGTGATGGACGACACCGTCAGTGCGTCCGTCGCGCAGGCTCTCGATGAGCTGATGCTTTGGGCCGAGGACTGCCCCGAAGTCCGG
CACCTCGTGCACGCGGATTTCGGCTCCAACAATGTCCTGACGGACAATGGCCGCATAACAGCGGTCATTGACTGGAGCGAGGCGATGTTCGGGGATTCCCAATACGAGGTCGCCAACAT
CTTCTTCTGGAGGCCGTGGTTGGCTTGTATGGAGCAGCAGACGCGCTACTTCGAGCGGAGGCATCCGGAGCTTGCAGGATCGCCGCGGCTCCGGGCGTATATGCTCCGCATTGGTCTTG
ACCAACTCTATCAGAGCTTGGTTGACGGCAATTTCGATGATGCAGCTTGGGCGCAGGGTCGATGCGACGCAATCGTCCGATCCGGAGCCGGGACTGTCGGGCGTACACAAATCGCCCG
CAGAAGCGCGGCCGTCTGGACCGATGGCTGTGTAGAAGTACTCGCCGATAGTGGAAACCGACGCCCCAGCACTCGTCCGAGGGCAAAGGAATAATcTCGAGTCATGTAATTAGTTATGTC
ACGCTTACATTCACGCCCTCCCCCCACATCCGCTCTAACCGAAAAGGAAGGAGTTAGACAACCTGAAGTCTAGGTCCCTATTTATTTTTTTATAGTTATGTTAGTATTAAGAACGTTATTTATATTT
CAAATTTTTCTTTTTTTTCTGTACAGACGCGTGTACGCATGTAACATTATACTGAAAACCTTGCTTGAGAAGGTTTTGGGACGCTCGAAGGCTTTAAcgagctcgaattcatcgatCAGTGTCGAAAACG
AGCTCGAATTCTTATTACGATCCTCGCGCCCCCTACCCACCGTACTCGTCAATTCCAAGGGCATCGGTAAACATCTGCTCAAACTCGAAGTCGGCCATATCCAGAGCGCCGTAGGGGGCG
GAGTCGTGGGGGGTAAATCCCGGACCCGGGGAATCCCCGTCCCCCAACATGTCCAGATCGAAATCGTCTAGCGCGTCGGCATGCGCCATCGCCACGTCCTCGCCGTCTAAGTGGAGC
TCGTCCCCCAGGCTGACATCGGTCGGGGGGGCCGTCGACAGTCTGCGCGTGTGTCCCGCGGGGAGAAAGGACAGGCGCGGAGCCGCCAGCCCCGCCTCTTCGGGGGCGTCGTCG
TCCGGGAGATCGAGCAGGCCCTCGATGGTAGACCCGTAATTGTTTTTCGTACGCGCGCGGCTGTACGCGGACCCACTTTCACATTTAAGTTGTTTTTCTAATCCGCATATGATCAATTCAAGG
CCGAATAAGAAGGCTGGCTCTGCACCTTGGTGATCAAATAATTCGATAGCTTGTCGTAATAATGGCGGCATACTATCAGTAGTAGGTGTTTCCCTTTCTTCTTTAGCGACTTGATGCTCTTGATC
TTCCAATACGCAACCTAAAGTAAAATGCCCCACAGCGCTGAGTGCATATAATGCATTCTCTAGTGAAAAACCTTGTTGGCATAAAAAGGCTAATTGATTTTCGAGAGTTTCATACTGTTTTTCTGT
AGGCCGTGTACCTAAATGTACTTTTGCTCCATCGCGATGACTTAGTAAAGCACATCTAAAACTTTTAGCGTTATTACGTAAAAAATCTTGCCAGCTTTCCCCTTCTAAAGGGCAAAAGTGAGTAT
GGTGCCTATCTAACATCTCAATGGCTAAGGCGTCGAGCAAAGCCCGCTTATTTTTTACATGCCAATACAATGTAGGCTGCTCTACACCTAGCTTCTGGGCGAGTTTACGGGTTGTTAAACCTTC
GATTCCGACCTCATTAAGCAGCTCTAATGCGCTGTTAATCACTTTACTTTTATCTAATCTAGACATATGAATTAATTCGGGCCGCGGAGGCTGGATCGGTCCCGGTGTCTTCTATGGAGGTCAA
AACAGCGTGGATGGCGTCTCCAGGCGATCTGACGGTTCACTAAACGAGCTCTGCTTATATAGACCTCCCACCGTACACGCCTACCGCCCATTTGCGTCAATGGGGCGGAGTTGTTACGAC
ATTTTGGAAAGTCCCGTTGATTTTGGTGCCAAAACAAACTCCCATTGACGTCAATGGGGTGGAGACTTGGAAATCCCCGTGAGTCAAACCGCTATCCACGCCCATTGATGTACTGCCAAAAC
CGCATCACCATGGTAATAGCGATGACTAATACGTAGATGTACTGCCAAGTAGGAAAGTCCCATAAGGTCATGTACTGGGCATAATGCCAGGCGGGCCATTTACCGTCATTGACGTCAATAGG
GGGCGTACTTGGCATATGATACACTTGATGTACTGCCAAGTGGGCAGTTTAGCGTAAATACTCCACCCATTGACGTCAATGGAAAGTCCCTATTGGCGTTACTATGGGAACATACGTCATTATT
GACGTCAATGGGCGGGGGTCGTTGGGCGGTCAGCCAGGCGGGCCATTTACCGTAAGTTATGTAACGCGGAACTCCATATATGGGCTATGAACTAATGACCCCGTAATTGATTACTATTAATA
ACTAGTCAATAATCAATGTCAACATGGCGGTAATGTTGGACATGAGCCAATATAAATGTACATATTATGATATGGATACAACGTATGCAATGGGCCAAGCTCCTCGAGTAATTCGCGCCACTTCT
AAATAAGCGAATTTCTTATGATTTATGATTTTTATTATTAAATAAGTTATAAAAAAAAATAAGTGTATACAAATTTTAAAGTGACTCTTAGGTTTTAAAACGAAAATTCTTGTTCTTGAGTAACTCTTTCCTGT
AGGTCAGGTTGCTTTCTCAGGTATAGCATGAGGTCGCTCTTATTGACCACACCTCTACCGGCAGATCAATTCCTCGATCGAGTTTACCACTCCCTATCAGTGATAGAGAAAAGTGAAAGTCGA
GTTTACCACTCCCTATCAGTGATAGAGAAAAGTGAAAGTCGAGTTTACCACTCCCTATCAGTGATAGAGAAAAGTGAAAGTCGAGTTTACCACTCCCTATCAGTGATAGAGAAAAGTGAAAGTC
GAGTTTACCACTCCCTATCAGTGATAGAGAAAAGTGAAAGTCGAGTTTACCACTCCCTATCAGTGATAGAGAAAAGTGAAAGTCGAGTTTACCACTCCCTATCAGTGATAGAGAAAAGTGAAAG
TCGAGCTCGGTACCCTTGTTGAGTTGTCCTCTCAGTTTTTATAAAAGTTTTTATTTTACCTTGTTTTGTAAATTAGATCTTTCAATTGGTTGGAATCATCTTAATTGAGTTATAGCAAACAGATCCACTA
GTGGCCTATATGGGTGCTAAAGAAAAGAGTGCTTGTCCAAAAGACCCTGCTAAACCTCCTGCCAAGGCTCAAGTTGTAGGATGGCCACCTGTTAGATCATACAGAAAGAATGTCATGGTTTCT
TGTCAAAAGTCTTCAGGTGGACCAGAAGCTGCAGCCTTTGTTAAAGTATCTATGGATGGTGCCCCATATTTGAGAAAGATTGACTTAAGAATGTACAAGggtgctggagcaggcATGGAGGAAGTGC
AAGACCTCGCAGCGGCGAGAGTCCGATCTCTGGCGAAAGTTAAGCGCATCTGGCACAATATCATTCAAAAATACTCCTCGCTGGGCGAGGATGAACAGGGTGATGTCTTGGATATGAGGA
CCGGAGACATTGTTCAGGATGTCGGCCATCTCCGATCAATGCGCGATTCCACCACCGTCAAAGGTGAGGTATGGAATAGCCTCATTGAACTGGAACGCAATAGAAGCCCCAAGAGTGAGA
AAATACGCCGGAAATCAGGAGACCTGATACCGTCGAGCCCTTCAAAAAAAGCAAGAAACTCCTGTGGTACATCCGCATTTACATCAATACTCATAGCAAGCGACTCTGAGGACGACGTGAC
GCAGTATTTTGACTCTTATAGGAGCTCGCCGCGTCGCACAACGCGATCCTCCATCGCCCAAAAGGACAATTTATCGGTTGTGCCGTCTCTCGATACGAACGCGTCTCCGCTTACGCCAATT
CGCAAAATGAGACGACCCAAATCCATTGGTGAAGACCCGCTACGCTTATCTCCACCAGCATACAATCTTGACGAAAACTACAAAATGTCGAGAAGATCGCTATTCAAGACCAAACCTGAAAA
TTCCGAGGGCCCGTTGATAACTCCATCCAAAGTGCGCCGGTTCAAGGCTGCGCTCGCACGTAAATCTGACGAGTCCGGGAACACTCGTTTGAAACGTCTGCTCTGGCTGAACAAACAAAA
AAAAGTGTGATTGCATCACAAAAACATAGGTGTCGCGCGACAATGCATTTGCACAGCCCTTGAGGCTAAAATAATTTCTAGAAAAACTGGGGACCTCAAAATAAATAGTCCCAATCGATTCTTT
ACGTTCAGACGAAAATGAGCCGCTCTCAGGTGTACTTTGACGTTTCCATTGGAGGTGAAACCGCAGGTCGTATTGTCTTCGAGCTTTTCAATGACATTGTTCCGAAGACGGCTGAGAACTTCA
GATCACTTGCGACGGGAGAACCAGGCTTTGGTTACCAGGGATCTATTTTCCATCGGGTGATCAAAAACTTTATGATCCAGGGTGGAGACTTCACTAATTTTGATGGCACGGGTGGCAAGTCAA
TATATGGTGAAAAGTTCGAAGACGAGAATTTCGAACTAAAGCATGACAAGCCGTTCTTACTTTCCATGGCCAATGCTGGAAAAAATACCAATGGCTCTCAGTTCTTTATCACCACAGTTCCAAC
ACCGCACCTAGATGGTAACCTGCATTAATGAATCGGCCAACGCGCGGGGAGAGGCGGTTTGCGTATTGGGCGCTCTTCCGCTTCCTCGCTCACTGACTCGCTGCGCTCGGTCGTTCGGC
TGCGGCGAGCGGTATCAGCTCACTCAAAGGCGGTAATACGGTTATCCACAGAATCAGGGGATAACGCAGGAAAGAACATGTGAGCAAAAGGCCAGCAAAAGGCCAGGAACCGTAAAAAG
GCCGCGTTGCTGGCGTTTTTCCATAGGCTCCGCCCCCCTGACGAGCATCACAAAAATCGACGCTCAAGTCAGAGGTGGCGAAACCCGACAGGACTATAAAGATACCAGGCGTTTCCCCC
TGGAAGCTCCCTCGTGCGCTCTCCTGTTCCGACCCTGCCGCTTACCGGATACCTGTCCGCCTTTCTCCCTTCGGGAAGCGTGGCGCTTTCTCAATGCTCACGCTGTAGGTATCTCAGTTCG
GTGTAGGTCGTTCGCTCCAAGCTGGGCTGTGTGCACGAACCCCCCGTTCAGCCCGACCGCTGCGCCTTATCCGGTAACTATCGTCTTGAGTCCAACCCGGTAAGACACGACTTATCGCC
ACTGGCAGCAGCCACTGGTAACAGGATTAGCAGAGCGAGGTATGTAGGCGGTGCTACAGAGTTCTTGAAGTGGTGGCCTAACTACGGCTACACTAGAAGGACAGTATTTGGTATCTGCGCT
CTGCTGAAGCCAGTTACCTTCGGAAAAAGAGTTGGTAGCTCTTGATCCGGCAAACAAACCACCGCTGGTAGCGGTGGTTTTTTTGTTTGCAAGCAGCAGATTACGCGCAGAAAAAAAGGATC
TCAAGAAGATCCTTTGATCTTTTCTACGGGGTCTGACGCTCAGTGGAACG
