## Supplemental Table 1 for "Centromere chromatin may regulate the Flip-flop type of the Mating Type Switching in methylotrophic yeast *Ogataea polymorpha*"

Supplementary Table S1. *O. polymorpha* strain list

| Strain | Genotype | Source |
|---|---|---|
| BY21401 | CBS4732 wild type | lab stock |
| HPH221 | NCYC495 *MATα* wild type | lab stock |
| HPH713 | NCYC495 *mat-a1Δ::hphNT1 Δku80::zeo ura3-1* | lab stock |
| HPH1622 | CBS4732 *rad6Δ::natNT2* | this study |
| HPH1746 | NCYC495 *IR1Δ::natNT2 ura3<<SLA2 Δku80::zeo FAS2-3'<<IR-URA3* | this study |
| HPH2089 | CBS4732 *ku80Δ::zeo ura3-1* | this study |
| HPH2099 | CBS4732 *ku80Δ::zeo ura3-1 ade12Δ::URA3-IR-IR-hphNT1* | this study |
| HPH2246 | NCYC495 *ku80Δ::zeo leu1-1<<LEU1-OsTIR-TetR'-Ssn6 ura3-1* | lab stock |
| HPH2272 | NCYC495 *3mCherry-dcu-ScLEU2fragment-natNT2-IR2 mata1Δ::hphNT1 ku80Δ::zeo ura3-1* | this study |
| HPH2273 | NCYC495 *IR2ΔmatαΔ-3mCherry-natNT2 FAS2-3'end::IR-OpKanMX ku80Δ::hphNT1 ura3-1* | this study |
| HPH2282 | NCYC495 *ku80Δ::zeo leu1-1<<LEU1-OsTIR-TetR'-Ssn6 ura3-1 iAID1-OpSCM3-hphNT1* | this study |
| HPH2286 | NCYC495 *ku80Δ::zeo leu1-1<<LEU1-OsTIR-TetR'-Ssn6 ura3-1 CNP1-AID-a-OpURA3* | this study |
| HPH2287 | NCYC495 *Δku80::zeo leu1-1<<LEU1-OsTIR-TetR'-Ssn6 ura3-1 CNP1-AID-OpURA3* | this study |
| HPH2329 | NCYC495 *ku80Δ::zeo leu1-1<<LEU1-OsTIR-TetR'-Ssn6 ura3-1 αΔ-natNT2* | this study |
| HPH2331 | NCYC495 *Δku80::zeo leu1-1<<LEU1-OsTIR-TetR'-Ssn6 ura3-1 CNP1-AID-OpURA3 αΔ-natNT2* | this study |
| HPH2332 | NCYC495 *ku80Δ::zeo leu1-1<<LEU1-OsTIR-TetR'-Ssn6 ura3-1 iAID-OpSCM3-hphNT1 Δalpha-natNT2* | this study |
| HPH2577 | CBS4732 *ku80Δ::zeo ura3-1 rad18Δ::hphNT1* | this study |
| HPH2599 | CBS4732 *bre1Δ::hphNT1 ku80Δ::zeo ura3-1* | this study |
| HPH2600 | CBS4732 *RAD6-5flag-hphNT1 ku80Δ::zeo ura3-1* | this study |
| HPH2601 | CBS4732 *RAD6ΔC-5flag-hphNT1 ku80Δ::zeo ura3-1* | this study |
| HPH2602 | CBS4732 *RAD6-GFP(ST)-hphNT1 ku80Δ::zeo ura3-1* | this study |
| HPH2603 | CBS4732 *RAD6ΔC-GFP(ST)-hphNT1 ku80Δ::zeo ura3-1* | this study |
