## Supplemental Table 2 for "Centromere chromatin may regulate the Flip-flop type of the Mating Type Switching in methylotrophic yeast *Ogataea polymorpha*"

Supplementary Table S2. Primer list

| Primer | Sequence |
|---|---|
| Hp_contig16_DIC1_Fw | TGTTGCCTGTCTTAGCAGCG |
| Hp_contig16_RV2 | ATAAGTACTCACAATCGAGGC |
| Hp_contig17_5 | AGGAACAGGTTCAGTACTGG |
| HpCNP1-S1 | TTATTTCCAAAGGGGTAATCAAAAAATTGTCCTGATAAACTTTAGGGCATTGTTTCAATGcgtacgctgcaggtcgac |
| HpCNP1-S2 | AAAAGGCCCGATTGGTATTTGACACAACAGGTTGTAGTATACATACACTAATAGGACTCAatcgatgaattcgagctcg |
| hph-Rv | CCCAAAGCATCAGCTCATCG |
| HpRAD18-S1 | CGCGTCGCGCAATCGAAAAATATTCAAATACTCTATTTGCAAGATCACTGATGGCGACATTCACTGATCCGCTCGACTCCcgtacgctgcaggtcgac |
| HpRAD18-S2 | CGCGAAAATGAGCATCGGCAGCATGCCGTACATCCCGTCGCTCAAGCTGGTGGAAAGTAAGGACATAATAGAATAGATTAatcgatgaattcgagctcg |
| HpRAD6_S1 | ATTTATATTTTAATAGCATATTCTTTTTCGTTCCATTGGCAAAAACTACGTTAGGAAATGcgtacgctgcaggtcgac |
| HpRAD6_S2 | AGGCGGTCAAATAGACATGACTTGTGCTCTCTAACGAACTATATGCTATTATGGTCGCTAatcgatgaattcgagctcg |
| HpURA3_16 | GCACAGAGAGACATGGGAGG |
| HpURA3_7 | cgtacgctgcaggtcgacTCCAGAACCTGCTACACCAT |
| MAT_38 | CGAAGTGGAAAACTGAGTGG |
| MAT-8 | Ttaaacaaggtagcaccgaaaaa |
| OpADE12_9 | ATTACACTATCGACGCGTGC |
| OpBRE1_S1 | GAAGGAGATACTCAGGTCCGGTTACGCTCATATTTCGCAAATATTGTATATTAGTTTGTCcgtacgctgcaggtcgac |
| OpBRE1_S2 | TCGATGCACGATAAACAATAATCCAAACTACATCATCAAGTTGGCATCGATTAGTACTTAatcgatgaattcgagctcg |
| OpRAD6_del-Ctail_S3 | TACAAAGACCACCGCAGCCAGTACATCAAACGAGTTAGACAGACTGTTGAGAAGAGCTGGcgtacgctgcaggtcgac |
| OpRAD6_S3 | GACGAAGACGACGAGGACGACGAGGACGGTATGGAAAAGGGAGAGGCGGACAACAAAGAGcgtacgctgcaggtcgac |
| OpSCM3_9 | ACCATCTAGGTGCGGTGTTG |
| OpSCM3_S1_AgTEF1 | TCGATCAGAATATTTTGGGATATTTTTTTATTTATTTTTCCCTTTATTTACATTTATATGgacatggaggcccagaatac |
| S2_TIR9myc_Rv | atcgatgaattcgagctcgGGaTCTCTtgaatgatcgttcc |
